# Managing risks and trade-offs in multispecies fisheries: the role of trophic control and price asymmetry

**DOI:** 10.1101/2025.01.29.635461

**Authors:** Théo Villain, Jean-Christophe Poggiale, Nicolas Loeuille

## Abstract

Ecosystem-based fisheries management (EBFM) aims to balance ecological and economic goals while employing precautionary measures to address uncertainties in knowledge and management. This study investigates fishing strategies to achieve this balance while minimizing the associated risks. Using a simplified prey-predator model, we explored various scenarios reflecting the diversity of global fisheries. The model integrates key ecological dynamics (bottom-up and top-down forces), economic factors (price structures), and aggregates the diversity of fishing practices by distributing fishing mortality across species. Our findings reveal that predators are always more affected by fishing than prey, regardless of the effort distribution. High yields are achieved by reducing predator densities—either maximizing predator catches when they are highly valued or reducing predation pressure to enhance prey harvests when prey prices are high. Such strategies result in significant ecological impacts, leading to systematic trade-offs. Elevated prey prices and top-down controlled systems intensify these trade-offs, increasing ecological risks. Regarding uncertainties, we demonstrate that maximizing yields poses risks to both biodiversity and profitability. Reconciliation is challenging but feasible when both species are fished. This balance can be achieved only in bottom-up controlled systems where prey valuation is not disproportionately high compared to predator prices.

## Introduction

Fishing is a critical activity for meeting global food needs, yet it significantly threatens the composition and future of marine communities. While the impacts of fishing are now well-documented, they affect all species—whether targeted or not—in an uneven manner (Daskalov et al., 2007; Ortuño Crespo and Dunn, 2017). For example, in 2022, the FAO estimated that 30% of fish stocks were exploited unsustainably (FAO, 2022), while a study covering 118 years of trawling presented an even more alarming picture, suggesting that 88% of species are overexploited, with biomass losses reaching up to 94% (Thurstan et al., 2010). On a global scale, for several decades, small fish at low trophic levels have accounted for an increasing share of total catches, giving rise to a phenomenon known as “Fishing down the marine food web” (Pauly et al., 1998). Nets are capturing fewer top predators, suggesting drastically reduced abundance (Christensen et al., 2014; Ferretti et al., 2008). It is estimated that 90% of large-bodied individuals, most often top predators, have disappeared from the oceans over the latter half of the 20th century (Myers and Worm, 2003).

Management policies have been implemented, such as those based on Maximum Sustainable Yield (MSY), but their relevance at the ecosystem scale has proven to be limited (Mackinson et al., 2009). Such criteria rely on single-species approaches to fishery productivity, at odds with larger scale objectives. When targeting small highly productive species, these policies suggest the extraction of a significant portion of their biomass (Hjermann et al., 2004; Pinsky et al., 2011; Smith et al., 2011). Resource depletions then significantly impact predator productivity, subsequently stabilizing prey densities, which further encourages such fishing practices. Conversely, neglecting predator control of prey in predator-targeted fisheries would lead to increased prey densities and cascading effects (e.g, (Estes and Palmisano, 1974; Gregr et al., 2020), ultimately impacting fishery profitability. These limitations have fueled growing interests in multispecies management strategies, based on the entire ecosystem (Ecosystem-Based Fisheries Management, EBFM, (White et al., 2012)).

EBFM aims to balance the preservation of marine community functioning with the interests of various stakeholders in a fishery (Pikitch et al., 2004). A key component lies in understanding the ecological mechanisms that regulate exploited systems. Trophic interactions shape ecosystem functioning and explain the high sensitivity of predators to fishing activities. Predators are generally large-bodied species (Woodward et al., 2005) with long generation times which makes them particularly vulnerable to exploitation (Jennings et al., 1999). Positioned at the top of food webs, they suffer not only from direct fishing pressures but also indirectly from fishing their prey (Ferretti et al., 2010). These dynamics can be more or less pronounced depending on the feedbacks within the network, making it crucial to understand ecosystem functioning in order to achieve sustainable exploitation.

Historically, species abundance at each trophic level was primarily thought to be regulated by resource availability. In systems controlled in such a bottom-up (BU) manner, predator biomass is highly constrained by prey densities (thick black arrows in “Ecosystem functioning” box Fig. 1). For example, fluctuations in plankton abundance constrain higher trophic levels and regulate the reproductive success of seabirds in the North Sea (Frederiksen et al., 2006), while marine food webs off the coast of Chile largely fluctuate with coastal upwellings, demonstrating the significance of this control (Cury et al., 2000). However, broader and more recent research has shown that top-down (TD) controls (ie, biomass being controlled mostly by upper trophic levels) occur in many marine ecosystems (Baum and Worm, 2009; Estes et al., 1978), as represented by thick red arrows in “Ecosystem functioning” box of Fig. 1. Large variations in the populations of mid-trophic fishes following the depletion or recovery of their predators highlight the structuring role of TD control (Baum and Worm, 2009). The relative frequency of top-down and bottom-up controls has been a subject of debate for many years but theoretically underpins the functioning and stability of ecosystems (Barbier and Loreau, 2019; Chase et al., 2000; Hunter and Price, 1992). It appears that there is spatial variability in how ecosystem dynamics are controlled, with upwelling regions being often BU controlled for instance (Baum and Worm, 2009). Similarly, some food webs are more resource regulated at the base and predator controlled at the top, with key intermediate species mediating the interaction between these two mechanisms (Cury et al., 2000; Lynam et al., 2017).

**Figure 1.**
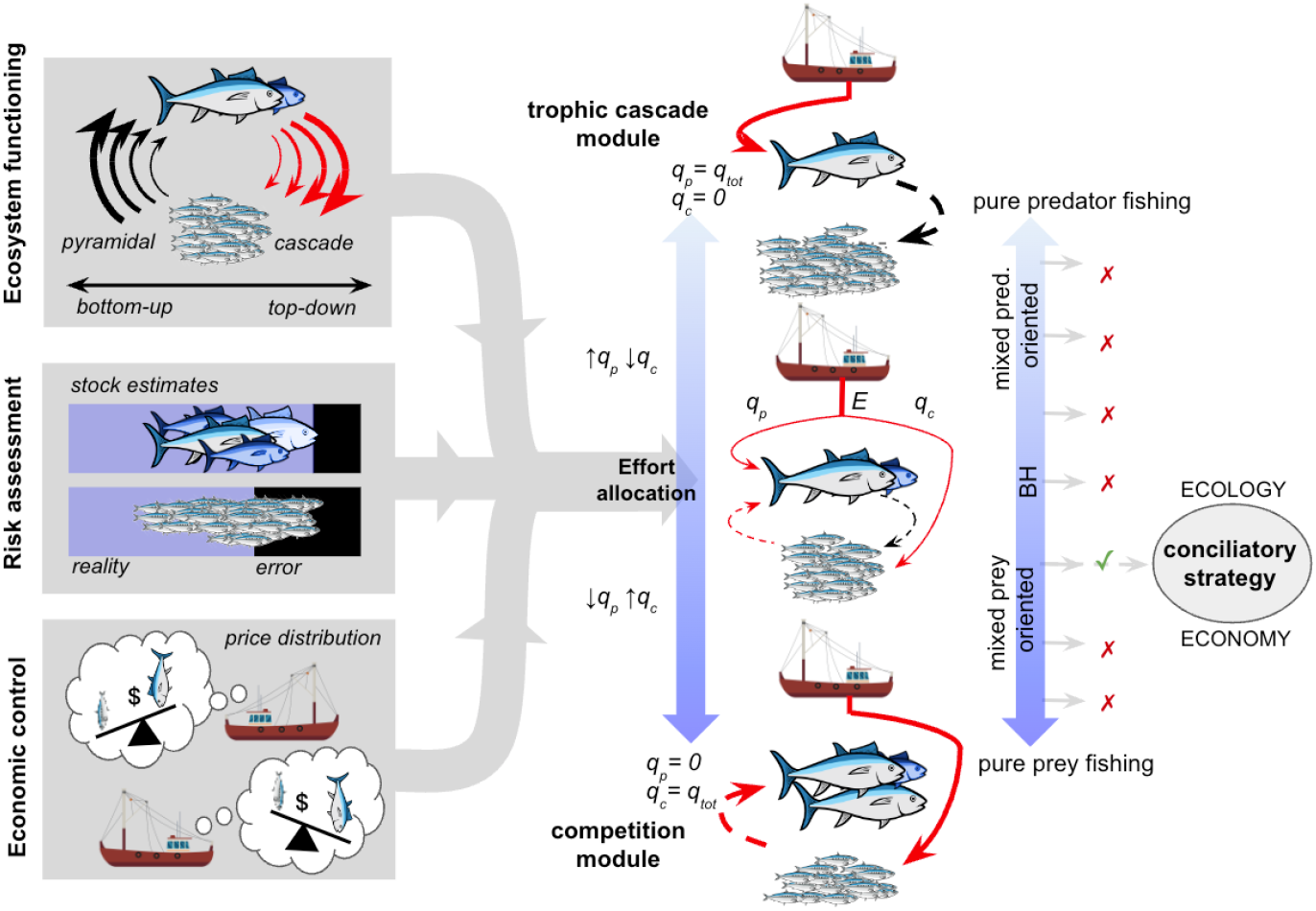
Designing the optimal fishing strategy from a socio-ecological perspective is constrained by the functioning of the ecosystem, the sensitivity to errors, and the economic structure of the fishery. Effort is allocated between prey and predator based on these dimensions. All scenarios are accounted for, from pure predator fishing when fishermen act as top-predators (trophic cascade module) to pure prey fishing where fishermen act as predator competitors (competition module). *q*_*i*_ : catchability of species *i*, and for a given effort, the fishing pressure on the ecosystem is constant (*q*_*c*_ + *q*_*p*_ = *q*_*tot*_). Black (red) arrows: positive (negative) effect on abundances. Solid (dashed) arrows: direct (indirect) effects.

The economic structure of fisheries is another key component of EBFM. Economic constraints can heavily influence the allocation of fishing quotas and shape catch patterns (“Economic control” box Fig. 1). The most profitable fish are generally the first to be targeted (Sethi et al., 2010), and since larger fishes tend to fetch higher prices, predators are often captured first (Pinnegar et al., 2002; Tsikliras and Polymeros, 2014). However, the development of aquaculture and the reduction of some forage fishing quotas have likely driven up the prices of small fishes (Villasante et al., 2013). Subsidies can also artificially increase the profitability of fisheries to the detriment of their ecological status (Munro and Sumaila, 2002). Accounting for economic incentives, while understanding how the ecosystem is regulated, is therefore essential for designing sustainable management policies.

Finally, scientific uncertainties (e.g. in abundance estimates, parameter calibration), as well as social uncertainties (e.g. in policy implementation and effectiveness), may endanger the efficiency of fisheries management (Fogarty et al., 1996; Privitera-Johnson and Punt, 2020). Taking these uncertainties into account, a retrospective analysis of abundance estimates for 230 fisheries worldwide revealed that the number of collapsed stocks were vastly overestimated at the time (1.85 fold, Edgar et al., 2024). These uncertainties must therefore be accounted for (Fig. 1, “Risk Assessment” box) when proposing a sustainable socio-ecological management approach for complex systems (Holsman et al., 2017).

In designing EBFM policies, new approaches suggest distributing fishing-induced mortalities across different trophic levels in proportion to their productivity (Garcia et al., 2012; Zhou et al., 2010). This distribution, known as balanced harvesting (BH), has been studied using both simple and complex models (Jacobsen et al., 2014; Plank, 2018; Zhou and Smith, 2017) and appears to potentially offer higher yields while maintaining the structure and functioning of ecosystems. However, these approaches have been criticized, particularly for their challenging field implementation (Froese et al., 2016; Pauly et al., 2016), and their study is generally based on the comparison of only a few alternatives (Zhou and Smith, 2017). A perfect balance, as much as a strong focus on a single species

(mono-specific management) is also a particular reference point. Here, we aim at considering all possible distributions of exploitation efforts between species, beyond particular reference points, to elucidate the best strategy in terms of ecological and economic conciliation. We particularly investigate how trophic control, price asymmetry, and uncertainties constrain the economic and ecological reconciliation of management policies. To provide a general response and an analytical understanding of underlying mechanisms, we developed a simplified model in which both prey and predator can be fished and in which these various aspects can be easily implemented.

We propose that among a range of strategies that sustain both species, those that most effectively limit predator densities in the ocean are likely to maximize economic returns, while a greater focus on prey is safer from an ecological perspective. This creates an inevitable trade-off between economy and ecology that ultimately determines the optimal distribution of effort. Indeed, from a profitability perspective, maximizing catches at higher trophic levels by targeting predators when they are most valuable, and reducing predation pressure when prey species hold higher value, present in both cases advantageous strategies. If these predictions are verified, restricting predator densities will have the greatest impact on ecosystem structure, as it significantly disrupts species density distributions. Strategies that exclusively target prey minimize the impact of fishing on biomass pyramids because they proportionally affect predator dynamics. Optimal distribution of fishing efforts between predators and prey will however likely change between BU and TD controlled systems. In BU systems where prey self-regulation controls system dynamics, capturing predators is expected to be less economically beneficial compared to TD-controlled systems. Consequently, we anticipate higher economic returns in TD systems compared to BU systems, especially when prey prices exceed those of predators. We believe that calibration errors are particularly detrimental to both economics and ecology when targeting predators since they are more sensitive to exploitation than prey. While prey-predator price asymmetry and interaction balance can economically influence optimal strategies, we expect that they do not alter the ecology/economics trade-off, suggesting that balanced fisheries might indeed benefit from reconciling these two objectives.

## Methods

To capture the vast diversity of global fisheries, we use a simplified socio-ecological model, enabling the analysis of numerous scenarios. From an ecological perspective, the diversity of exploited species can be summarized into two groups: prey and predators. This classification is based on the ecological effects of fishing (Christensen, 1996): fishers may either compete with predators for their prey (competition module, Fig. 1) or target predators, thereby benefiting prey populations (cascade module, Fig. 1). Either outcome depends on the relative importance of BU and TD forces, which are critical for understanding ecosystem functioning implications of fishing (Barbier and Loreau, 2019). From an economic perspective, the diversity of fisheries stakeholders, fishing campaigns, and techniques can be aggregated into a shared pool of effort, which impacts the entire ecosystem. The allocation of this effort between prey and predators reflects different management strategies. This simplified view of the system’s ecology (Lotka-Volterra prey-predator model), exploitation (capturability and effort), and economics (multispecies economic yield) provides a mathematical framework to understand the underlying mechanisms. While the model we use is arguably highly simplified, it allows a systematic investigation of the key components of the question, and its simplicity allows us to provide a mathematical analysis of optimal strategies, and numerical investigations whose robustness is simpler to assess.

We use a Lotka-Volterra model with two compartments in trophic interaction, and the mortality associated with fishing is distributed proportionally to their production (eq. 1 & 2):

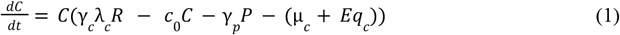

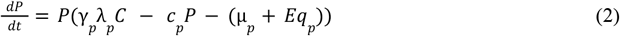

*C* and *P* are the respective densities of prey (consumer) and predator, *γ*_*i*_, *λ*_*i*_, μ_*i*_ and *q*_*i*_ the respective attack rate, conversion efficiency, intrinsic mortality rate and fishing catchability of species *i. R* is the resource density, *c*_0_ (*c*_*p*_) the per capita intraspecific competition rate of prey (predator) species and *E* the fishing effort. All parameter values are detailed in Supplement 1 with ecological parameters calibrated using size-based relationships (Brose et al., 2006; Brown et al., 2004), see Supplement 2). Because our aim is to understand how the partitioning of fishing effort affects the conciliation of economy and ecology, we fix total catchability and allow prey and predator catchabilities to vary, so that *q*_*tot*_ = *q*_*c*_ + *q*_*p*_, with *q*_*c*_ and *q*_*p*_ the respective catchabilities of preys and predators. We consider the entire continuum of strategies in this work, whose names correspond to the scale shown in Fig. 1.

In the first part, we mathematically explore how fishing effort and catchability influence abundances and characterize trophic control within such a system.

In the second part, we examine the conciliation of economic and ecological aspects, considering all possible fishing strategies by varying effort and catchability independently, without preconceived notions about the effects of such manipulations from an application perspective. Economic returns are assessed based on the multispecies economic yield (eq. 3) to understand the impact of economic incentives in a multispecies context. Defining this yield does not require knowledge about the operational aspects, as only the overall fishing effort *E* is considered.

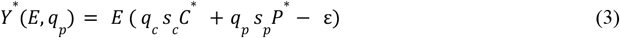

Prey and predator respective catchabilities distribute the fishing effort on prey or predator populations. The profitabilities of the two populations therefore depend on their relative abundances and on their respective per individual selling prices *s*_*c*_ and *s*_*p*_ (or *s*_*ik*_ per biomass unit of species *i*), with a cost ε proportional to the effort involved (Dichmont et al., 2010). Impacts of fishing strategies on ecology are captured based on diversity measures. The Shannon diversity index (eq. 4) accounts for the maintenance of the two species as well as for their relative frequency.

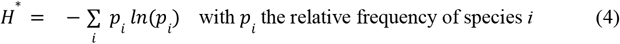

While this index is most usually applied in complex, highly diverse communities, we here use it because it embodies not only variations of species diversity, but can also be perceived as a proxy of maximum trophic height or of the pyramid of abundances. Indeed, values of *exp*(*H*^*^) here by definition go from 1 to 2. Values close to 1 indicate that the system does contain predators, but at very low density (hence a low trophic height and a very bottom-heavy pyramid), while values close to 2 would indicate that predators and prey are equally abundant (higher trophic height and more homogeneous pyramid). We aim to understand how the type of trophic control and price asymmetry influence these impacts.

In the third part, following traditional approaches, we assume that fisheries management follows the multispecies maximum sustainable yield (MMEY, e.g. (Hoshino et al., 2018)). We compute this criterion for different distribution of efforts, and compare how these various MMEYs satisfy the conciliation of economic and ecological objectives. This implicitly assumes, in contrast to the second part, that controlling fishing effort (days at sea) is easier than controlling prey catchability (type of technology used on vessels). For a given distribution (pool of technologies), there exists an effort 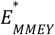 that maximizes profits in the long term:

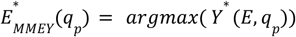

This effort depends on the fishing strategy, and thus the different yields and Shannon index at MMEY are given by:

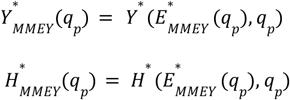

We evaluate the effect of managing at 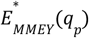 on the ecology/economy trade-off and also discuss the degree of risk associated with MMEY strategies, given uncertainties in stock assessments (Edgar et al., 2024) and in management implementation. This analysis covers a wide range of systems, considering both trophic control and market price structure.

In the final section, we aim to propose strategies that effectively reconcile ecology and economics by introducing thresholds related to management measures effectiveness. In practice, we will evaluate ecologically 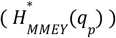 and economically 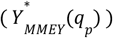 all strategies within a given socio-ecological system and retain those that exceed 60% of the maximum values obtained.

## Results

When exploitation pressure is very important, fishing leads to the extinction of both species. This extinction equilibrium occurs whenever 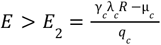 (see Supplement 3). As intuitively expected, pure predator-oriented strategies (*q*_*p*_ = *q*_*tot*_ ie. *q*_*c*_ = 0) cannot lead to prey extinction.

Below this level of pressure, prey first occur alone at a density 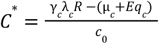. This prey-only equilibrium is stable whenever the exploitation does not allow predators persistence 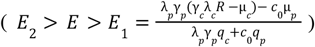 (see Supplement 3). Both species coexist once the effort is lower than *E*_1_ . Under this condition:

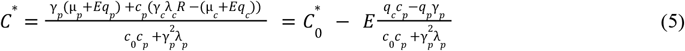

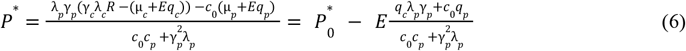

Where 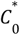 and 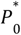 are consumer and predator density without exploitation respectively. From eq. 6, note that predators suffer from fishing in two ways: direct fishing *q*_*p*_, amplified by prey self-regulation *c*_0_ and indirect effects of fishing prey, amplified by predation pressure *λ*_*p*_*γ*_*p*_. The relative magnitude of these direct and indirect effects on predator abundances reflects the importance of BU forces (*c*_0_) and TD forces 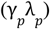.

### 1 Opposite or parallel variations of species, depending on ecological conditions

Effects of exploitation on densities can lead to simultaneous decreases of all species or to opposite variations, depending on predator catchability:

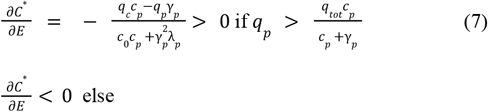

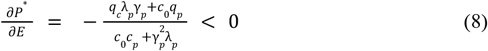

Predators indeed always suffer from higher effort (eq. 8) while prey benefit when predator catchability is high 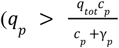, eq. 7) due to reduced predation. Similarly, biasing efforts toward predators increases all densities in TD controlled systems (eq. 10), but harms predators when the system is BU controlled.

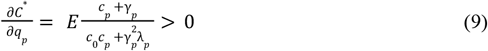

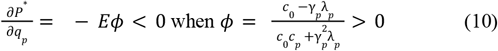

Prey indeed consistently benefit from a greater focus on predator fishing (eq. 9) because they are less exploited (*q*_*c*_ = *q*_*tot*_ − *q*_*p*_) and face reduced predation pressure. When *ϕ* is positive (*c*_0_ > *γ*_*p*_ *λ*_*p*_, ie dominant BU controls (Barbier & Loreau 2019)), predators suffer from concentrated fishing efforts targeting them 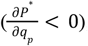,even though their prey are less fished and thus more abundant. In this scenario, the impact of fishing on predators is primarily determined by prey self-competition and productivity, rather than predator’s capacity to transfer energy. As a result, reducing fishing pressure on low-productivity prey does not offset the fishing over-mortality of predators. Conversely, when *ϕ* is negative (dominant TD controls), predators benefit from fishing efforts being concentrated on them rather than on their prey (eq. 10). In this case, predators are more limited by their capacity to acquire energy than by prey productivity (*c*_0_ > *γ*_*p*_ *λ*_*p*_). Fishing less on highly productive prey increases prey densities, which in turn benefits predator populations due to high trophic transfers. The trophic control term (*ϕ*) therefore encapsulates in a continuous manner how focusing exploitation on predators affects their density through TD (*γ*_*p*_ *λ*_*p*_) and BU forces (*c*_0_). These equations highlight how changes in the ecological context (trophic controls) alter fishery-induced feedbacks and species sensitivity to fishing. Based on this, increasing effort is always detrimental ecologically as predators are eventually lost. Balanced strategies seem preferable for BU controlled systems, while a more predator oriented exploitation allows better ecological states in TD controlled systems.

### 2 A difficult conciliation of economic and ecological objectives

Ecological and economic objectives are difficult to reconcile. However, our results suggest that a prey oriented fishing (here *q*_*p*_ about 0.1) and moderate efforts (here around 20 effort units) offer a narrow range of strategies allowing conciliation (Fig. 2). This reconciliation is quite hard because optimal ecological states require low efforts and/or low predator catchabilities (green on Fig. 2), coexistence being rapidly lost in other conditions (gray on Fig. 2), while economic objectives typically require higher efforts and predator catchabilities (yellow on Fig. 2). Predators are more sensitive to exploitation than preys (mathematical analysis above), thus maintaining a complete ecological system requires maintaining predators and limiting density imbalances (eq. 4). Since effort and catchability combination determine the direct mortality from fishing (− *q*_*i*_ x *E*), maintaining predators at high efforts can only be achieved by mainly targeting prey (high *q*_*c*_ / low *q*_*p*_, e.g., red dashed line Fig. 2A). Reducing effort means that predator fishing can be increased (convex form of the Shannon index on Fig. 2, white and dark green areas) without leading to their disappearance (gray area). If community structure is less damaged by fishing when targeting preys (Shannon index approaching its maximal value when *q*_*c*_ = *q*_*tot*_ on Fig. 2B-C-D), reducing effort leads to less steep Shannon-index variations and allows for exploring more fishing strategies including balanced distributions (Fig. 2B-C-D).

**Figure 2.**
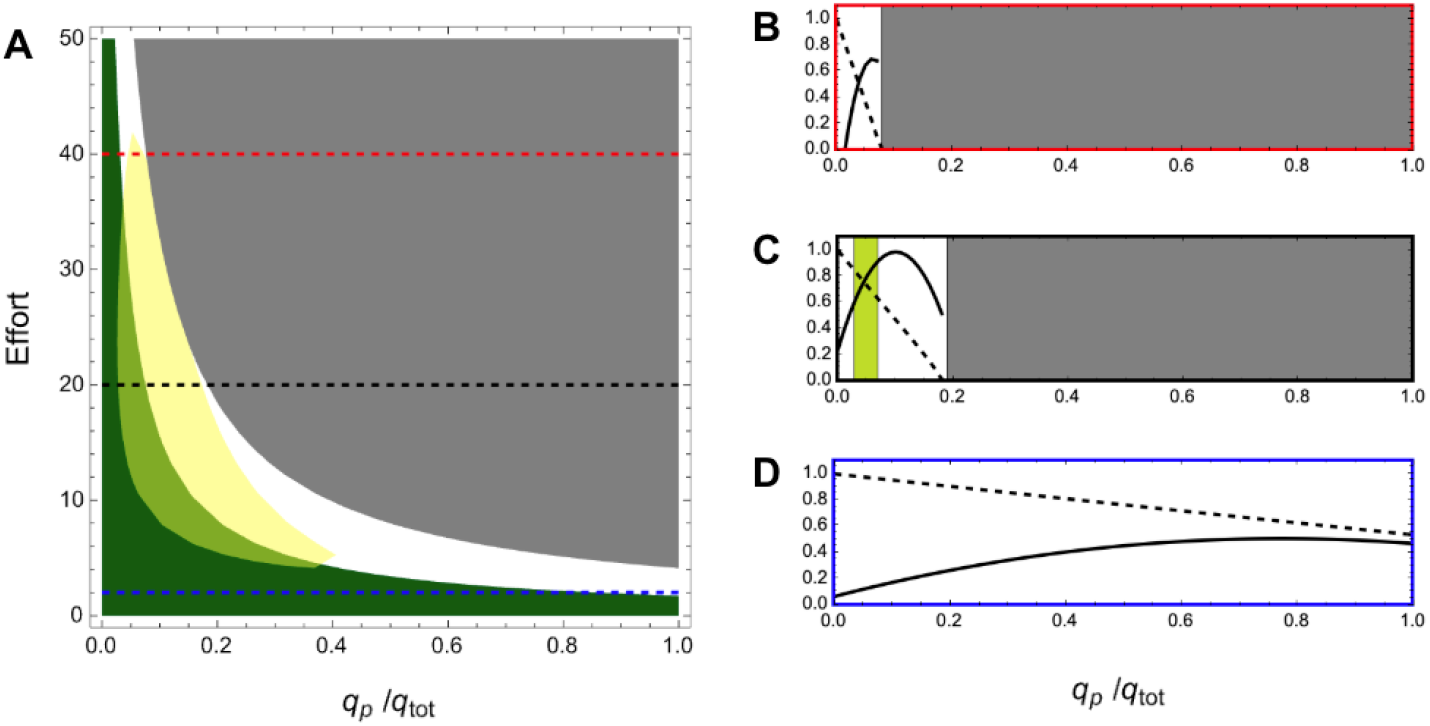
Conciliation of ecological and economic objectives along different efforts and fishing strategies. Objectives are considered achieved if fishing yields or Shannon indexes exceed 60% of their respective maximum values. A: Fisheries meeting economic (ecological) objectives are highlighted in yellow (dark green). Dashed lines represent transects of constant effort, detailed in B-C-D with respective border colors. B-C-D: Normalized Shannon index (dashed) and economic yield (solid) as functions of fishing strategy under constant effort. Light green areas: optimal strategies where economic and ecological objectives are simultaneously achieved. Gray areas in all panels indicate predator extinction. Price asymmetry: *s*_*pk*_ = 7, *s*_*ck*_ = 5. Trophic control parameters set to default values.

The multispecies economic return (*Y*^*^) depends on catches of prey, predators and their relative prices (eq. 3, Fig. 2, Fig. S4). When predator price exceeds prey price, as commonly observed (and as implemented on Fig. 2), predator fishing is valued so that maintaining healthy predator populations is especially important. Note however that predators are rapidly lost if fishing intensity is increased or exploitation more predator focused. This sets an upper limit to predator oriented fishing, in spite of their intrinsic value, leading to the counter-intuitive result that optimal strategies are largely oriented towards prey in spite of predator high prices. Prey fishing only is unattractive because it prevents benefiting from the market value of predators and because there is no beneficial effect from the removal of predation on prey.

Importantly, the position of the conciliation strategies depends both on economic (e.g., price structure, Fig. S4) and on ecological aspects (e.g., BU or TD forces, Fig. S5). Concerning economy, yield can be maximized by strategies targeting predators exclusively when specializing solely in prey is loss-making 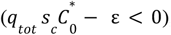, for instance because prey prices are very small (Fig. S4A). In this case, predator catches need to be maximized, with a great fishing focus on their stocks and therefore low effort (*E* = 1. 2 Fig. S4D). This situation allows a slightly larger set of conciliation strategies (Fig. S4A), oriented towards predators. Conversely, when prey price exceeds predator price, conciliatory strategies are hard to find (though a small set exists at high effort (*E* ≃ 40) and focusing on prey capture (Fig. S4E)). Along a predator-prey market value ratio, as it becomes increasingly attractive to maximize prey catches, the optimal strategies are shifting from a mixed predator-oriented/pure predator fishing with low effort to mixed prey-oriented fisheries heavily exploited (Fig. S4E). Note that BH is not necessarily adapted to all economic constraints since it leads to conciliation only when prey fishing is loss-making (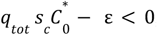, Fig. S4C).

On the ecological side, the value and sign of *ϕ* (BU/TD) influences the density ratios (eq. 10) and therefore modifies the response of the Shannon-index to catches. The increase in TD control (lower *ϕ*) does not have the same implications for management depending on whether it arises from a decrease in BU forces or an increase in TD forces, due to their opposing effects on densities. Indeed, conciliatory strategies become increasingly prey-oriented when prey self-regulation (BU forces) is reduced (Fig. S5A), while they shift their focus towards predators as predation pressure (TD forces) intensifies (Fig. S5B). Lower intraspecific competition increases prey and predator densities (eq. 5 & 6), but slightly changes density ratios (and therefore the global shape of the Shannon) since densities are qualitatively affected in a similar way. The overall fish biomass is increased which makes the system less sensitive to catches (higher Shannon-index) and leads to increased yields (eq. 3). Conversely, increased predation homogenizes prey-predator density ratios as changes in fishing then exert opposite effects on predators and prey densities (eq. 5 & 6).

Multiple combinations in BU/TD forces allow a given strength of trophic control *ϕ* (eq. 10). In a BU system (*ϕ* > 0), increasing TD control through the combined effects of weakened BU forces and strengthened TD forces leads to opposite conciliatory patterns depending on price asymmetries (Fig. S5). Optimal strategies are becoming increasingly predator-oriented when prey prices dominate (Fig. S5C), but increasingly prey-oriented in the opposite scenario (Fig. S5D). Increased TD control amplifies the benefits of reducing predation pressure on prey densities. When prey are highly valued, this encourages greater exploitation of predators to enhance possible prey benefits. Conversely, when predators are highly valued, it allows for increased exploitation of prey due to their higher abundance. Note, however, that TD systems (*ϕ* < 0) do not allow for objective reconciliation. In such systems, fishing predators release predation pressure to such an extent that driving predators close to extinction becomes economically advantageous, leading to strategies that are ecologically unsustainable.

### 3 Maximizing economic returns through MMEY management: deleterious effects on diversity and risks

While conciliation of ecology and economy remains an important objective, most fisheries are managed using an economy-oriented index (MSY or MEY). We therefore now leave out the conciliation side, assume that the system is exploited based on MMEY, and investigate the consequences of such a choice. Figure 3 illustrates the combined influence of predator and prey prices on the strategy that maximizes profitability, as well as the resulting socio-ecological correlation. When economic yields are used to calibrate the optimum fishing effort 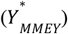, optimal fishing strategies are sensitive to the prey-predator price structure (Fig. 3A), but economic and ecological objectives are always anti-correlated regardless of the price structure (Fig. 3B-E showing all decreasing curves). Note however that the slope of the curve on these panels shows the ecological cost of increased yields, steeper slopes suggesting less costs. The comparison of panels B to E highlights that a focus on yields is especially harmful from an ecological point of view when prey is more valuable.

**Figure 3.**
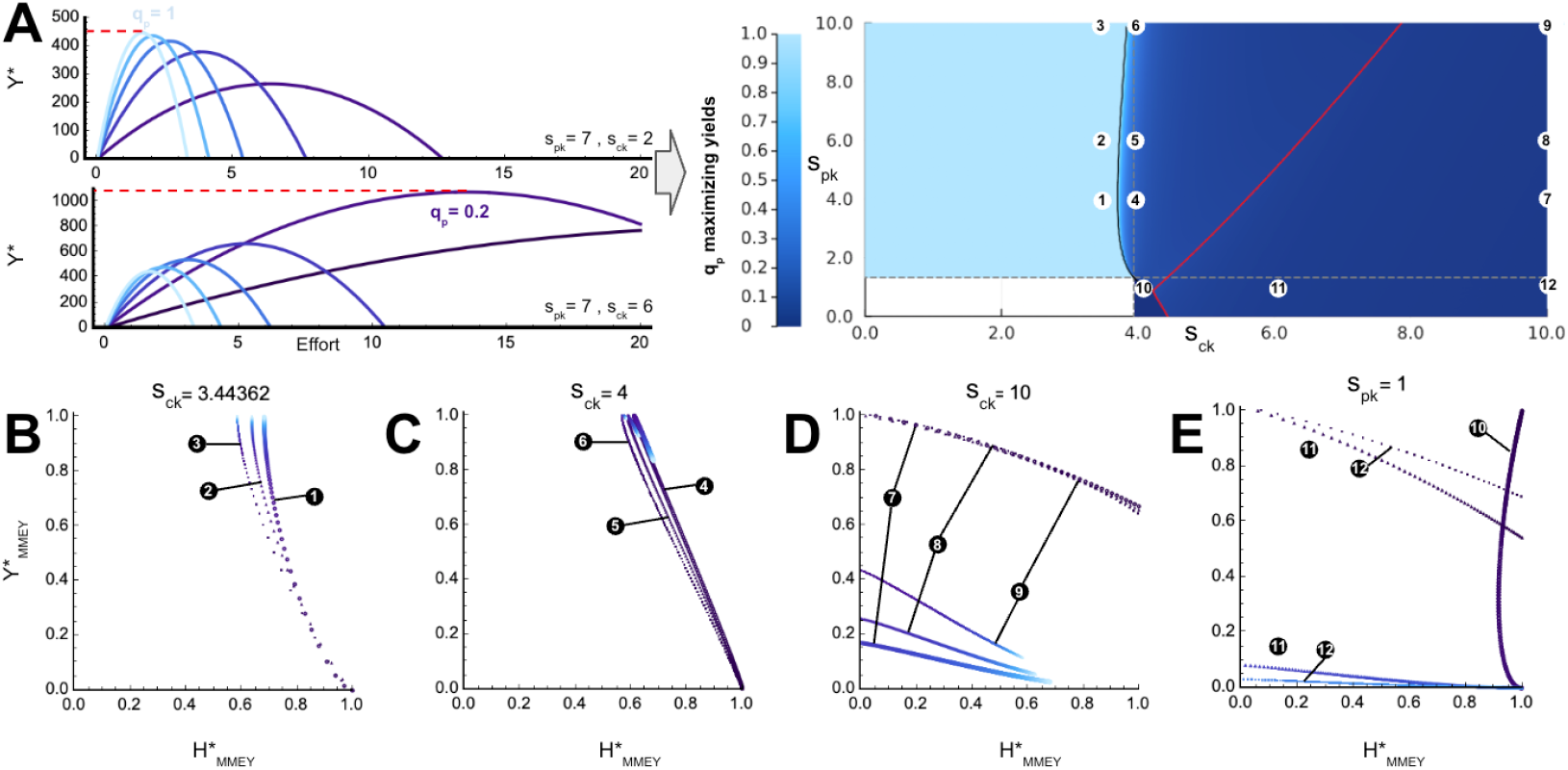
Impact of price structure (*s*_*ck*_, *s*_*pk*_) on strategies maximizing yield (A) and associated trade-offs (B-C-D-E). For a given price structure, the strategy (*q*_*p*_ /*q*_*tot*_) which maximizes profits among all possible strategies is depicted on panel A (right). Dashed lines: price structures for which monospecific predator (horizontal) or prey (vertical) fishing is unprofitable. Black line: prey yield contribution is significant compared to predator. Red-bordered area: price structures for which maximizing profits leads to predator extinction. B-C-D-E: Impacts of optimizing economic yields on 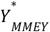 and on the resulting 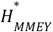 (normalized). Dot colors and numbers indicate the strategy and the associated price structure (panel A). Trophic control with default value.

Indeed, when comparing various market conditions (*s*_*ck*_, *s*_*pk*_), three distinct fishing strategies emerge: targeting only predators (light blue, Fig. 3A), engaging in mixed fisheries with a focus on prey (medium blue, Fig. 3A), or exclusively targeting prey (dark blue, Fig. 3A). We can show that the type of exploitation depends on combinations between three criteria:

i. whether such monospecific fisheries are economically sustainable or not:

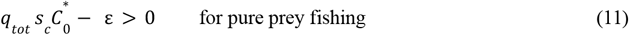

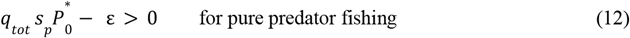
ii. whether pure predator fishing (eq. 13) or pure prey fishing lead to shortfalls (eq. 14) compared to other MMEY strategies (see mathematical developments in Supplement 10 eq. 14 & 15).

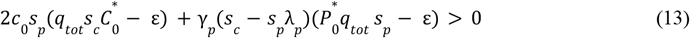

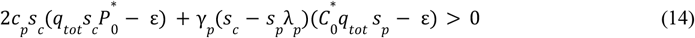
iii. whether the most profitable strategy would lead to predator extinction:

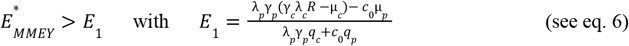

These limits are depicted by gray dashed lines (for (i)), by a black line (for (ii)) and by a red line (for (iii)) on Fig. 3A.

When the exploitation of predators alone is profitable (eq. 12) and prey contribute little to the MMEY, predator fisheries maximize yields (i.e.,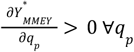, see Supplement 10) by maximizing predator catches. For a given price structure, these fisheries consistently minimize predator densities and consequently exhibit the lowest Shannon index (Fig. 3B). Note that prey contribution can be positive even when 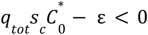 since mixed fisheries generally lead to 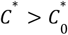 with preys benefiting from predator fishing (eq. 7).

However, the most profitable strategy shifts as soon as preys become sufficiently valued (*s*_*c*_) relative to predators (*s*_*p*_):

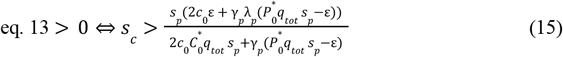

In this case, the most profitable strategies involve the exploitation of both species (as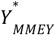 becomes a modal function of *q*_*p*_), although the exploitation is heavily prey-biased. It is advantageous to target both prey and predators due to their intrinsic market values, but also because reducing predation pressure enhances prey stocks. Since the strategy is prey-oriented, the fishing effort is high, which significantly reduces predator densities and consequently degrades the ecological quality (Fig. 3C). When the valuation of prey is even higher, fishermen earn more profit from extinguishing predators (Fig. 3D-E), which is catastrophic for the ecosystem. Finally, a last scenario may arise where maximizing profits involves exclusively targeting prey. If fishing predators does not result in direct profitability (eq. 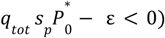 and prey prices are too low to justify predator removal, the selected fisheries are those that solely target prey (dark blue in Fig. 3A and curve 10 in Fig. 3E). However these conditions are not highly representative of the typical price ratios between prey and predators.

When BU control is weaker (lower positive *ϕ* Fig. S6) or even when the system is fully TD-controlled (*ϕ* < 0 Fig. S7), the ecology-economy trade-off is still verified. Maximizing returns in MMEY management is always to the detriment of diversity (Fig. S6 and S7 B-C-D-E). Price structures that maximize yields by capturing only predators are more restricted because prey densities are higher with lower *ϕ*. Wide price ranges allow for maximizing returns through mixed strategies but most lead to predator extinction (Fig. S6 red line and Fig. S7 red and yellow lines).

It becomes clear that the most profitable strategy for a given market condition depends on the trade-off between the direct and indirect value provided by predators, which is regulated by the value of prey. Increasing TD control leads to increased predator densities but their indirect contribution is even higher so that optimal economic strategies favor their extinction. Except when predator prices are so low that fishing predators is per se unprofitable, all the most profitable strategies maximize yields by minimizing predator densities, which explains the negative correlation between the two objectives.

Accounting for sources of uncertainty, maximizing economic returns from MMEY-managed fisheries is also very risky (Fig. 4), regardless of price structure and trophic controls (Fig. S8&S9). Figure 4 shows how departure from the optimal economic strategy (red line) affects economic and ecological aspects (from left to right) under different price ratios (from top to bottom). The most profitable strategy is to maximize catches of predators by only fishing them when their price greatly exceeds that of prey (red line in Fig. 4B-C). In this case, any error in estimating *E*_*MMEY*_ has an even greater impact on yields and diversity, as this is the strategy that leaves the fewest residual predators in the ocean. When prey price exceeds predator price, the most profitable strategy tends towards a state of predator exclusion (Fig. 3A&D-E and Fig. 4H-I) so any error in estimating *E*_*MMEY*_ leads to very large variations in profitability and diversity. When the prices of prey and predators are close, the most profitable strategy requires coexistence to take advantage of the market values of both species and reduce predation pressure (Fig. 3A&C and Fig. 4E-F). In this case, the densities of both species are low, and any estimation error strongly affects predator densities and therefore diversity. Note that in all cases, the optimal strategy from an economic point of view (red line) is maximally risky (or close to) compared to other strategies. Note also that underestimating *E*_*MMEY*_ always favor diversity (solid lines in Fig. 4C-F-I). When the BU control is weaker (lower positive *ϕ* Fig. S8), or even when the system is fully TD controlled (*ϕ* < 0 Fig. S9), returns are all the greater, but maximizing profits is once again the riskiest strategy and leads to predator extinction in the vast majority of cases.

**Figure 4.**
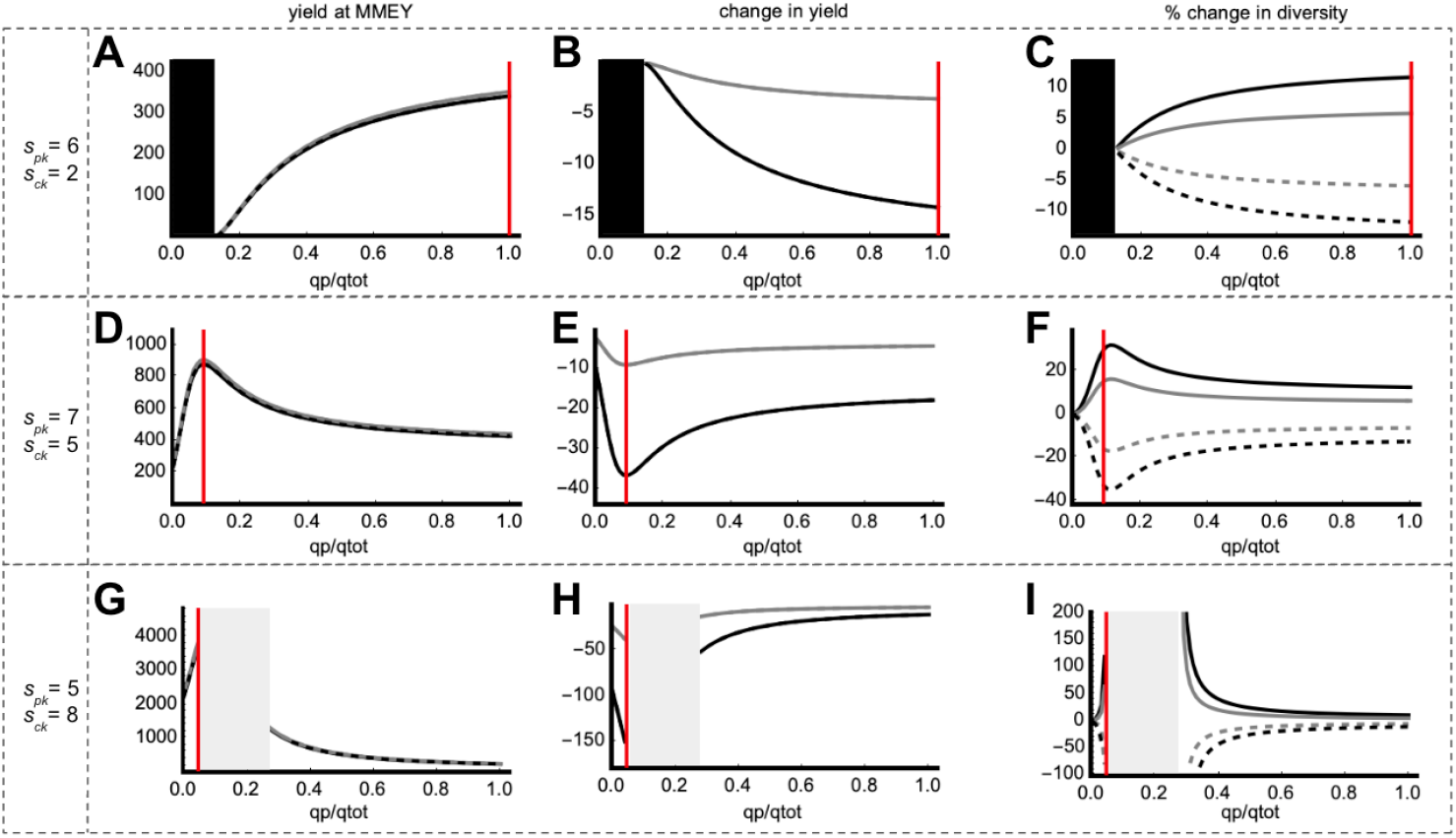
Impact of estimation errors in *E*_*MMEY*_ calibration on economic yield and diversity. Black (gray) lines: error of 20% (10%) in *E*_*MMEY*_ calibration. Solid (dashed) lines: underestimation (overestimation). The red vertical line depicts the fishing strategy producing the best economic yield. A-D-G: economic yields with calibration errors. B-E-H: effects of calibration errors on yields. C-F-I: effects of calibration errors on diversity. Price structure values are indicated on the left side of each line. Black areas: unprofitable fishing strategies. Gray areas: fishing strategies leading to predator extinction when maximizing yields with MMEY metrics (calculated without calibration errors). Trophic control with default value.

### 4 Socio-ecological conciliation through MMEY management: higher proportion of prey harvesting in mixed strategies

Whatever the price structure, regulating fisheries with MMEY in TD-controlled systems (*ϕ* < 0) does not allow for reconciliation of socio-ecological objectives since it is always extremely beneficial to suppress predation pressure for economic reasons. Figure 5 shows how the price structure (*s*_*pk*_ /*s*_*ck*_) and the trophic control (*ϕ*) impact conciliation strategies. In BU-controlled systems (*ϕ* > 0), socio-ecological conciliation occurs for mixed-fishing strategies (Fig. 5). These strategies consist of targeting mainly prey and less predators than the one that optimizes economic returns in MMEY management strategies (Fig. 6). More valuable prey lead to a reconciliation at lower predator capturabilities, for all trophic controls. This can be explained by the increase of prey price which weakens the effect on yield of predator captures and the lifting of predation pressure. Since the best distribution of effort on diversity objectives consists of fishing only for prey, the strategy that reconciles economic and ecological criteria is in between (Fig. 6). However, when the prices of the two species are similar, two ecological-economic conciliation zones emerge (Fig. 5 within the * area). This transition occurs when fishers, aiming to maximize their returns, shift from a fishing regime focused on maximizing predator catches to a regime that benefits from the intrinsic value of both species (eq 13 & eq 14). In this scenario, on either side of the most economically profitable strategy, it is possible to reconcile both objectives. When the price of prey becomes even higher, conciliation strategies increasingly prioritize prey fishing.

**Figure 5.**
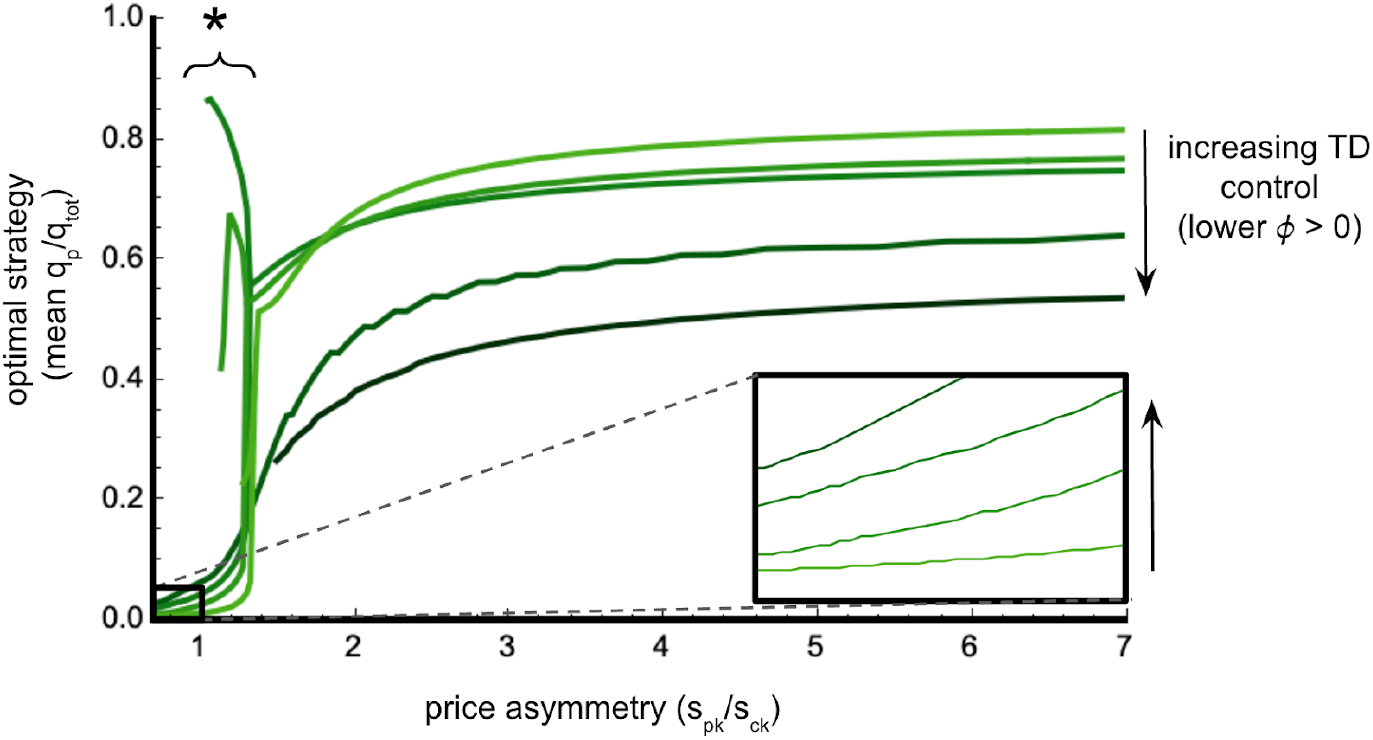
Conciliation strategies in MMEY management for different price structures (*s*_*pk*_ /*s*_*ck*_) and trophic controls (*ϕ*). For each socio-ecosystem structure, a range of strategies (*q*_*p*_ /*q*_*tot*_) conciliate economy and ecology if they exceed 60% of the most profitable strategy yield 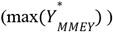 and of the best ecological status 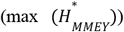. The mean strategy of the range is then reported. Two discontinued ranges of optimal strategies may exist, especially when prey-predator prices are close (* area). Colors depict different *ϕ* values, from light green (strong BU, high positive *ϕ*) to dark green (low BU, low positive *ϕ*). TD-controlled systems (*ϕ* < 0) do not allow any conciliation since it exacerbates the socio-ecological trade-off.

**Figure 6.**
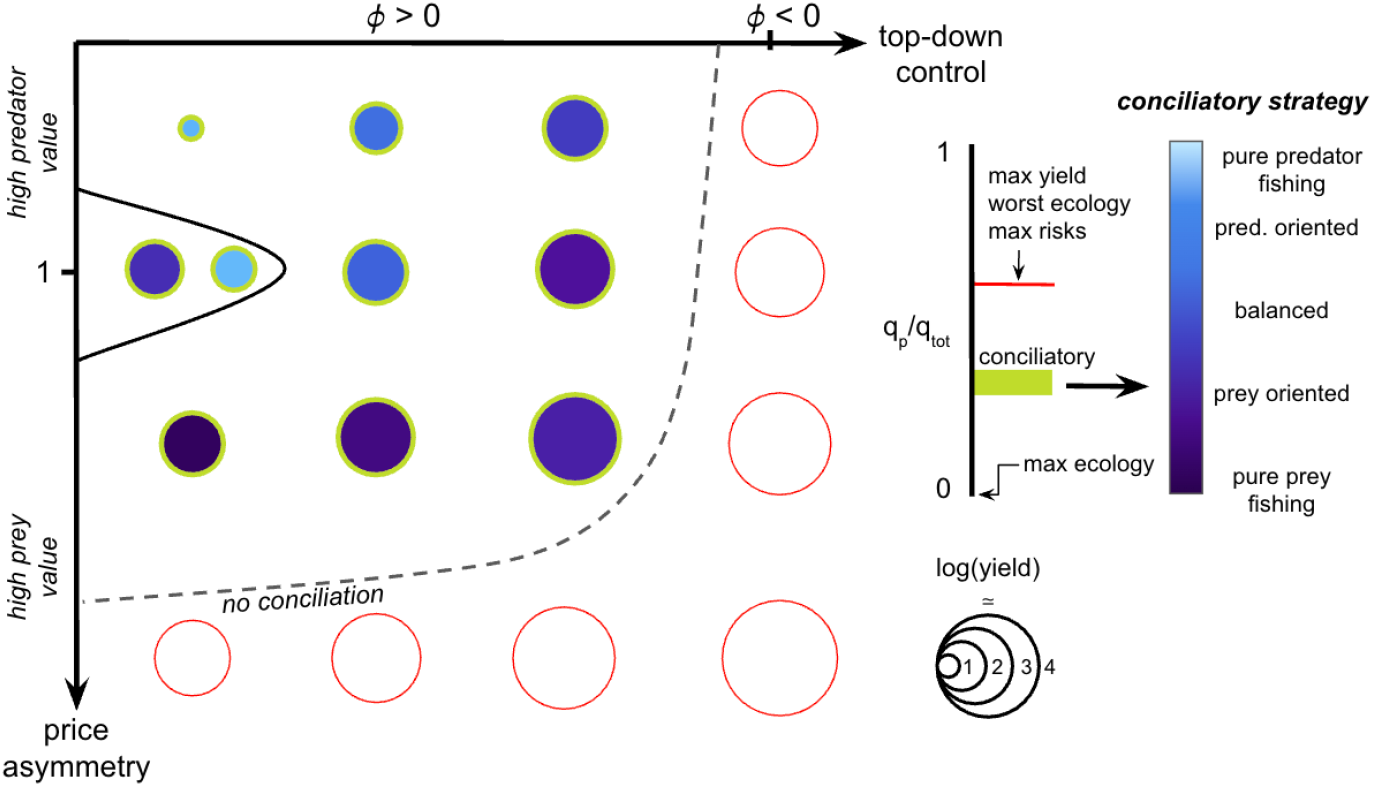
Conciliation of socio-ecological objectives is hard to achieve and only in bottom-up controlled ecosystems where prey prices are not too high compared to predator prices. This is achieved through mixed fishing strategies generally using an effort distribution between the most profitable fishing strategy (red line) and pure prey fishing. When predator and prey valuations are close, two conciliatory strategies may exist, on either side of the most profitable strategy. In bottom-up systems, increasing top-down control or prey prices makes conciliatory strategies more focused on prey. Circle sizes represent the yield of conciliatory (green border) or MMEY (red border) strategies if no conciliation is possible.

Finally, when TD control increases (BU forces decrease while TD forces increase) in a BU system (*ϕ* > 0), the effect on the conciliation strategy depends on the price structure in a counterintuitive manner, whereas no conciliation is possible in TD systems (*ϕ* < 0). When prey are valued higher than predators (Fig. 5, zoomed area), fishers primarily target prey for their high intrinsic value but also harvest predators to relieve predation pressure on prey. Increasing TD control amplifies the effect of predator removal on increased prey stocks. This explains why the conciliation strategy becomes slightly more oriented toward predator fishing, even if it is close to pure prey fishing. Conversely, when predators are more valuable than prey, fishers aim to maximize predator catches. Under increased TD control, removing prey has less impact on predator densities, thereby allowing greater prey fishing.

## Discussion

Our study reveals that reconciling ecological and economic objectives is often hard to achieve as inevitable trade-offs tend to emerge from the underlying ecosystem dynamics. When it is achievable, conciliation occurs through a mixed distribution of fishing effort (Fig. 2 & Fig. 5). These strategies generally lie between the most economically profitable distribution of effort and exclusively targeting prey (Fig. 6). Using metrics at MMEY to maximize yields makes fishing both economically and ecologically risky, thus supporting the implementation of intermediate strategies to balance socio-ecological objectives (Fig. 4). This mixed reconciliation reflects the anti-correlation between two criteria: fishing has less impact when targeting prey alone, while maximizing yields requires fishing predators. The price structure and trophic control shape the valuation of stocks and their responses to fishing (Fig. 3). We identified a *ϕ* parameter, which captures the effect of fishing on predators through the distribution of effort and ecological constraints. When predators are limited by prey self-competition, the system is resource-controlled and BU (*ϕ* > 0). As prey competition decreases and predators become more efficient in resource acquisition, optimal strategies become more balanced because reducing predation pressure becomes increasingly advantageous. Conversely, in TD systems (*ϕ* < 0), predator resource acquisition determines their densities more than prey self-competition. Allocating an increasing share of effort towards predators then benefits them by increasing prey abundance. Since predation pressure here determines prey densities, the effect of its release is greatly amplified which significantly penalizes predator-prey density ratios. In this scenario, we showed that ecological and economic reconciliation is not possible (Figure 5).

We demonstrated that predators are more heavily impacted by fishing than their prey, regardless of the effort distribution across compartments. This result aligns with current observations of predator depletion in the oceans (Myers and Worm, 2003; Pauly et al., 1998) and large predators sensitivity to fishing (Ferretti et al., 2010). While this is a direct consequence of trophic level positioning it also implies that, under constant effort, fishing prey is the most beneficial strategy for maintaining diversity, as it has proportional effects on predator density. This also shapes the ecosystem response to fishing, resulting in a convex Shannon index curve. At high fishing effort, diversity can be preserved if the effort is primarily directed toward prey, whereas at low effort levels, all strategies maintain an acceptable level of diversity. These results are consistent with empirical observations of sustainability in low-effort artisanal fisheries (Jul-Larsen, 2003) and consistent with previous theoretical analyses (Kolding et al., 2016; Tromeur and Loeuille, 2017). Therefore from a conciliatory perspective, higher prey prices favor fishing efforts in these systems but higher predator value decreases the overall effort.

We also showed that regardless of the trophic or price structure, the allocation of effort which produces the best MMEY inevitably results in minimizing predator densities. This limitation arises from two key phenomena. Predator species are larger, more expensive per unit of biomass (Sumaila et al., 2007; Swartz et al., 2013), rarer than their prey, and increasingly in demand (Willis and Bailey, 2020). Thus, when predators are highly valued, maximizing predator catches results in a significant reduction in their densities. Conversely, when prey are highly valued, predator densities are indirectly threatened by the fishermen’s tendency to substantially decrease predation pressure since both are competing for resource availability (Christensen, 1996; Hjermann et al., 2004). Therefore, irrespective of the socio-economic system, maximizing yields always comes at the expense of biodiversity, as commonly described (Cheung and Sumaila, 2008; Moore et al., 2021). However, highly valued prey stocks exacerbate this trade-off, making it even more costly for diversity and can drive predator species to extinction. These results highlight how the use of MMEY can threaten species with low productivity and economic value (Clark, 2010; Tromeur and Doyen, 2019) but also that management plans trying to maintain sufficient predator populations on the long run can be more resilient (Tromeur and Loeuille, 2017).

Finally, any errors in stock assessments used to compute the MMEY in estimating species interactions, or in implementing fishing strategies could lead to significant costs across ecological and economic dimensions (Pinsky et al., 2011). For instance, tunas are highly migratory, complicating stock estimation and management (FAO, 2022), with only 7 out of 15 commercially fished species being accurately assessed. Similarly, small pelagic fisheries are not protected from overexploitation (Pinsky et al., 2011) and are highly sensitive to climatic variability (FAO, 2022), with ocean warming exacerbating uncertainties in stock estimation. Thus, we demonstrated that regardless of the trophic or price structure, fishing becomes increasingly risky as the effort repartition produces higher MMEY. For a given fleet, calibrating the effort required to achieve maximum yields has been shown to be risky (e.g., Thorpe et al., 2017) and reducing it preserves bio-economic sustainability (Tromeur et al., 2021). Here, our study generalizes these results and emphasizes that among various techniques, the one possibly yielding the highest profits should also exhibit the greatest precision in fishing. The risk-yield relationship emerges here as a general pattern and highlights the necessity of adopting precautionary approaches to fisheries management (Lauck et al., 1998; Sethi, 2010).

As a consequence of these three findings, and as a general outcome of this study, it is possible to reconcile economic and ecological aspects while mitigating risks by adopting an intermediate distribution of fishing effort—generally between the effort allocation that maximizes yields and the one that targets only prey species. This result aligns with calls to reduce the high pressure on a few commercially targeted species and to distribute fishing-induced mortality across a broader range of species (balanced harvesting, Garcia et al., 2012; Zhou et al., 2010). Balanced fisheries have demonstrated promising outcomes when fishing-induced mortality is allocated proportionally to the natural productivity of species, as evidenced by studies employing complex models (Bundy et al., 2005; Jacobsen et al., 2014; Kolding et al., 2016) and simpler frameworks (Plank, 2018; Zhou and Smith, 2017). However, this concept has faced substantial criticism, particularly regarding the challenges of accurately estimating productivity and its practical implementation (Froese et al., 2016; Pauly et al., 2016). These critiques motivated the alternative approach adopted here, where overall fishing effort is distributed between two compartments proportionally to each species’ production. Nonetheless, our results align with the conclusions of these studies, as they advocate for simultaneously fishing across different compartments to reconcile economic and environmental objectives.

Price structure and ecosystem functioning jointly influence the socio-economic consequences of fisheries. First, higher prey prices lead to optimal strategies oriented on fishing prey, and the reverse applies to predator prices. This finding aligns with the observation that the first species to be heavily fished are those with the highest market value (Sethi et al., 2010). Notably, fishing exclusively for predators can be the most profitable strategy, regardless of whether fishing exclusively for prey is profitable or not. Conversely, pure prey fishing is rarely the most profitable strategy. This occurs only when predator fishing is itself unprofitable and fails to compensate for the gain in prey stock valuation driven by reduced predation pressure.

We found that the trophic control index (*ϕ*) plays a crucial role in defining fishing strategies. This result is not surprising, as consumer versus resource control shapes both the diversity and functioning of ecosystems (Worm et al., 2002) and determines their stability (Barbier and Loreau, 2019). First, the trophic control index illustrates how direct (fishing predators) and indirect (fishing prey) effects are amplified on predator densities. It depends on BU and TD forces highlighting the feedback mechanisms induced by fishing. Given the critical role of fishers’ positioning relative to predator dynamics (competition or cascade modules, Fig. 1), the index effectively reflects how the impacts of fishing propagate throughout the ecosystem. When positive, fishing has an overall BU effect, whereas a negative *ϕ* indicates a global TD effect (where predators benefit directly from predator-specific fishing). We found that in a BU system, weaker BU forces (lower positive trophic control *ϕ*) lead to a conciliation based on prey-oriented fishing. This is because prey benefit more from a weak BU effect than predators do, given their lower self-competition. Conversely, stronger top-down forces make predator fishing more advantageous for reconciliation, as this benefits predators at the expense of prey. However, when *ϕ* diminishes (a more TD-controlled system, with jointly reduced BU and increased TD forces), reconciliation strategies tend to become more balanced. This shift is due to the increased impact of predator fishing on prey densities. As a result, when prey are highly valued, this favors higher predator fishing to benefit on the resulting increased prey populations. Conversely, when predators are highly valued, it drives greater prey fishing, as their populations become abundant. These findings highlight two key points: failing to account for economic incentives in fishing limits the analysis of fisheries’ socio-ecological efficiency, and the system’s intrinsic BU and TD forces fundamentally shape its response to fishing.

We here use a simple, highly stylized model. Its simple structure still accounts for the three axes at hand (trophic control, price asymmetry and risk assessment), but it inevitably leaves out some economic and ecological complexities. From an economic perspective, the analyses conducted here assume equilibrium conditions, which obscure the economic challenges that multiple stakeholders may face during the implementation of fishing regimes. While optimal patterns capture long-term profitability for fisheries, transient effects are left out. Given our network context, prey and predator densities may undergo significant oscillations modulated by exploitation, potentially undermining the profitability and, consequently, the success of the fishing regime. Fluctuations in fish abundance can also lead to changes in global market prices, creating transient feedbacks. Transitional-phase analyses are therefore needed to complement the equilibrium-based conclusions.

From an ecological perspective, our simple approach does not capture the intrinsic complexity of ecological networks. While our approach provides a direct assessment of BU and TD effects, we leave out the multiple, direct and indirect interactions of varying intensities that occur in complex networks. Furthermore, we examined the effects of fishing primarily through changes in species abundances, whereas previous studies have shown that fishing can drive rapid eco-evolutionary dynamics (Olsen et al., 2004). In particular, the selectivity of fishing may lead to increased selection of certain phenotypes. For instance, increasing fishing selectivity on size at the species level to protect juveniles or benefit from large individuals may select slower growth phenotypes (Grift et al., 2003; Kuparinen and Merilä, 2007; Olsen et al., 2004) and reduce species production, potentially altering the conclusions of this study.

In spite of these caveats, we still offer some key messages that could be important from a management perspective. Particularly, we point out that trade-offs between ecology and economy are often very hard to overcome, conciliation requiring the fine-tuning of efforts and selectivity, on a narrow range. We also stress that some fisheries can be especially difficult to manage (e.g., when prey prices are high and the system is top-down controlled). While it may be possible to develop levers to alleviate economic constraints (e.g., adjusting prices through subsidies), intrinsic ecological vulnerabilities are much harder to overcome and our knowledge of the frequency of top-down vs bottom up controls is very often lacking. We hope that our model will encourage a proper consideration and empirical investigations of these key aspects.

## Supplementary

### Supplement 1: Table of parameters, units and default values

**Table S1:** Parameters used in the model. Species interaction parameters have been calibrated using size-based relationships (Brose et al. 2006, Brown et al. 2004 see Supplement 2). (1) Default values.Variations in these parameters are investigated to allow variations in trophic controls and price asymmetries.

| Parameters | Biological meaning | Unit | Value |
| --- | --- | --- | --- |
| $\gamma_c$ | Per capita attack rate of preys | $\text{ind}^{-1}\text{t}^{-1}$ | 10 |
| $\gamma_p$ | Per capita attack rate of predators | $\text{ind}^{-1}\text{t}^{-1}$ | $10^{(1)}$ |
| $\lambda_c$ | Conversion efficiency of preys | - | $2.87E - 3$ |
| $\lambda_p$ | Conversion efficiency of predators | - | $2.87E - 3$ |
| $\mu_c$ | Mortality rate of preys | $\text{t}^{-1}$ | $2.44E - 2$ |
| $\mu_p$ | Mortality rate of predators | $\text{t}^{-1}$ | $7.62E - 3$ |
| $c_0$ | Per capita intraspecific competition rate of preys | $\text{ind}^{-1}\text{t}^{-1}$ | $1^{(1)}$ |
| $c_p$ | Per capita intraspecific competition rate of predators | $\text{ind}^{-1}\text{t}^{-1}$ | 1 |
| $q_{tot}$ | Total catchability | $\text{t}^{-1}\text{effort}^{-1}$ | 1 |
| $q_c$ | Prey fishing catchability | $\text{t}^{-1}\text{effort}^{-1}$ | Range $[0, q_{tot}]$ |
| $q_p$ | Predator fishing catchability | $\text{t}^{-1}\text{effort}^{-1}$ | Range $[0, q_{tot}]$ |
| $s_c$ | Per capita prey selling price | $\$ \text{ind}^{-1}$ | $s_{ck} * e^{z_c}/1000$ |
| $s_p$ | Per capita predator selling price | $\$ \text{ind}^{-1}$ | $s_{pk} * e^{z_p}/1000$ |
| $s_{ck}$ | Per biomass prey selling price | $\$ \text{kg}^{-1}$ | $5^{(1)}$ |
| $s_{pk}$ | Per biomass predator selling price | $\$ \text{kg}^{-1}$ | $7^{(1)}$ |
| $\epsilon$ | Cost of effort | $\$ \text{effort}^{-1}$ | 125 |
| R | Basal resource density | ind | 5000 |

### Supplement 2: Life-history and interaction parameters

The life-history parameters and the strength of predatory interactions depend on relative prey and predator sizes (expressed as a log(biomass) *z*_*i*_). Mortality rate is assumed to be decreasing with size (Brown et al. 2004):

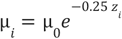

Attack rate is a function of predator-prey size ratio with a maximal value when predators are d times larger than their prey (Brose et al. 2006) :

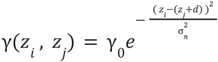 with *z*_*i*_ and *z*_*j*_ the respective predator and prey size.

The conversion efficiency is the efficiency of converting one unit of prey *j* into one unit of predator *i* and decreases with size difference:

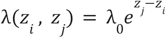

In this study, we consider ressource size *z*_*R*_ = 1, consumer size *z*_*c*_ = 5. 65, predator size *z*_*p*_ = 10. 3 and we assume preferred predator-prey size difference *d* = 4. 65 to be the same for consumers and predators. Competition is taken into account but is intraspecific and therefore does not depend on size differences (*c*_0_ = *c*_*p*_ = 1). Other parameters values: μ_0_ = 0. 1, *γ*_0_ = 10, *λ*_0_ = 0. 3, σ_*n*_ = 1.

### Supplement 3: Equilibrium feasibility and stability

The fishing intensity *E* and the distribution of effort between the two species (*q*_*p*_, *q*_*c*_ with *q*_*p*_ + *q*_*c*_ = *q*_*tot*_) determine the persistence of populations under exploitation. Specifically, the Lotka-Volterra dynamic system (eq. 1, eq. 2) exhibits three distinct equilibria (*C* *, *P* *):

*Eq*_1_ = (0, 0) where both species go extinct due to fishing pressure,

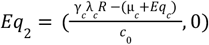 where predators are driven to extinction, while prey persist,

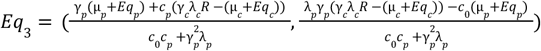 where fishing allows the coexistence of both species.

To analyze the feasibility and stability conditions of these equilibria, we define 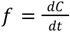 and 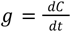 as the rate of change for prey and predator populations, respectively (eq. 1, eq. 2). The associated Jacobian matrix *J* is expressed as:

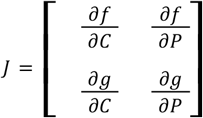

For each equilibrium, the Jacobian matrix *J* takes the following form:

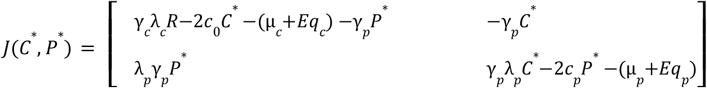

### Analysis of equilibrium *Eq*_3_

We aim to determine the fishing conditions that allow the coexistence of prey and predators. For *Eq*_3_ to exist, both population densities must remain positive. Prey populations persist as long as *E* < *E*_*c*_,
where:

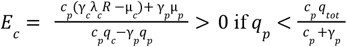

This critical threshold *E*_*c*_ represents the maximum fishing intensity beyond which prey populations cannot be sustained. Additionally, predator persistence requires that their population density remains above zero, which imposes further constraints on fishing strategies and effort distribution. When the effort distribution toward predators exceeds 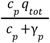, prey populations cannot go extinct. Predators, however, are maintained as long as *E* < *E*_1_, where:

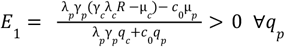

The difference *E*_*c*_ − *E*_*p*_ is positive if 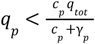. Specifically:

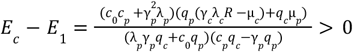

Thus, when *E* < *E*_1_, both prey and predators coexist in the system, making *Eq*_3_ feasible. To assess the stability of *Eq*_3_, we analyze the trace (*tr*) and determinant (*det*) of the Jacobian matrix *J* at equilibrium *Eq*_3_ .

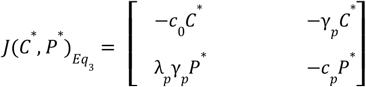

when and *P*\* > 0 *C*\* > 0

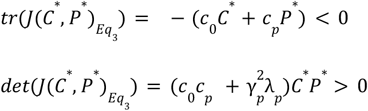

Therefore, when *P*\* > 0 and *C*\* > 0, in other words when *E* < *E*_1_, *Eq*_3_is feasible and stable.

### Analysis of equilibrium *Eq*_2_

The fishery maintains only prey populations if the fishing effort does not exceed *E*_2_ . Thus, when *E* < *E*_2_, *Eq*_2_ is feasible.

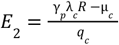

To assess the stability of this equilibrium, we examine the trace and determinant of the Jacobian matrix *J* at equilibrium *Eq*_2_:

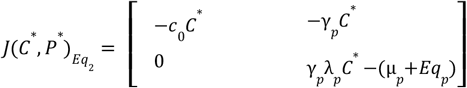

When *E* < *E*_2_, *C*^*^ > 0

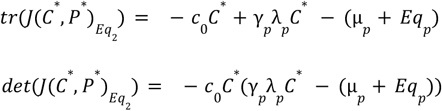

For *Eq* to be stable, the trace and determinant of *J* at *Eq* must be negative and positive, respectively. The feasibility conditions alone are not sufficient for the stability of *Eq*_2_. It is observed that:

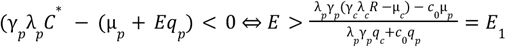

Therefore, when *E*_1_ < *E* < *E*_2_, *Eq* is stable.

### Analysis of equilibrium *Eq* _1_

*Eq*_1_ = (0, 0) and is always feasible.

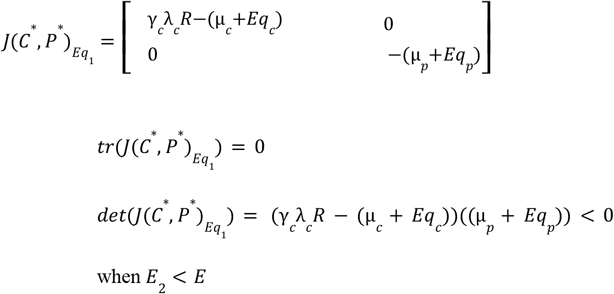

when *E*_2_ < *E*

Thus, when the fishing effort exceeds *E*_2_, the system reaches *Eq*_1_ (a saddle point).

### Supplement 4: Price distribution impacts on fishing strategies and objectives conciliation

**Figure S4.**
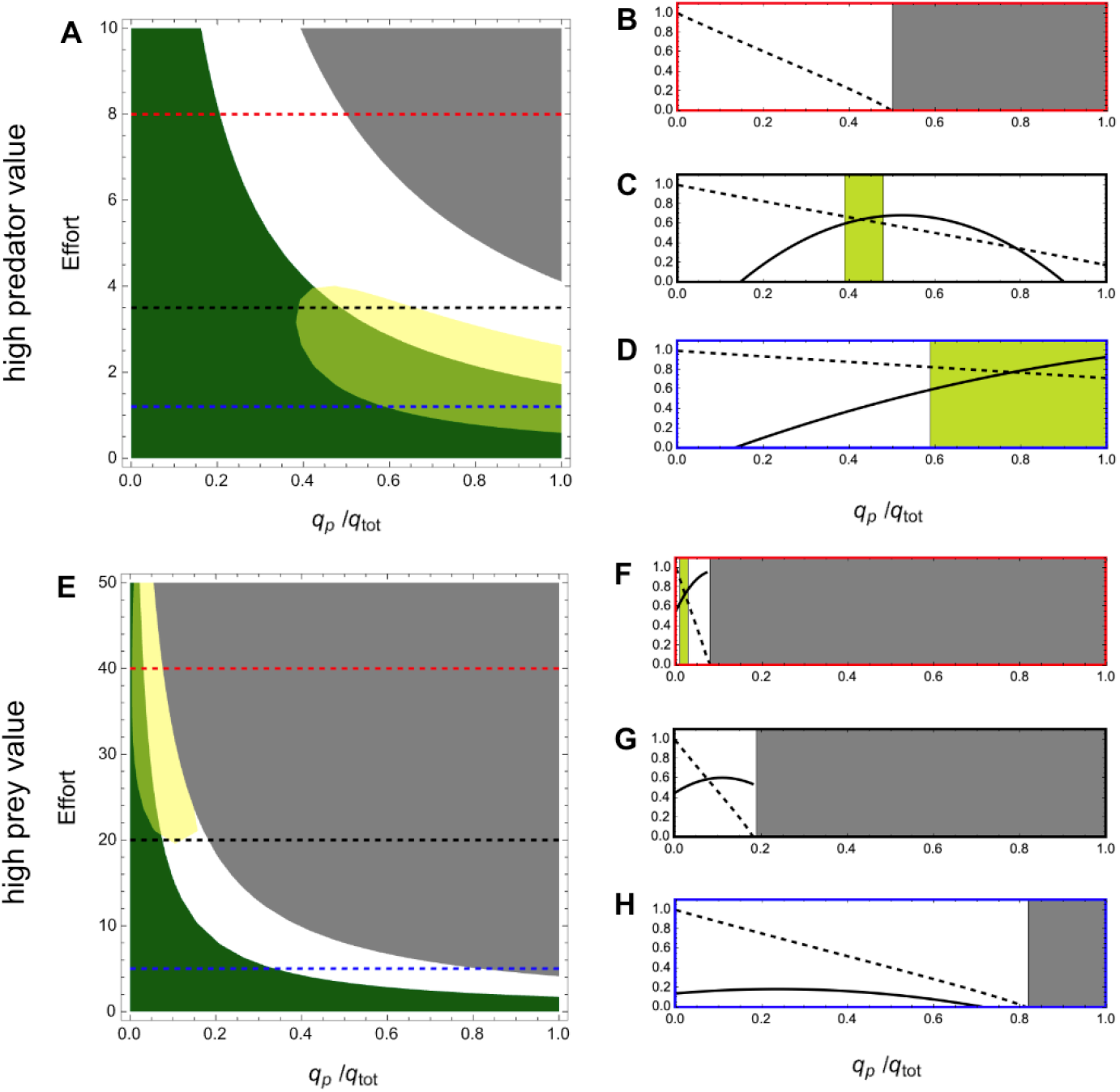
Impacts of price distributions on reconciling objectives. Objectives are considered achieved if fishing yields or Shannon indexes exceed 60% of their respective maximum values. Price asymmetry: *s*_*pk*_ = 6, *s*_*ck*_ = 2 (A) and *s*_*pk*_ = 5, *s*_*ck*_= 8 (E). A&E Fisheries meeting economic (ecological) objectives are highlighted in yellow (dark green). Dashed lines represent transects of constant effort, detailed on their respective right sides with the same border colors. B-C-D & F-G-H: Normalized Shannon index (dashed) and economic yield (solid) as functions of fishing strategy under constant effort. Light green areas: optimal strategies where economic and ecological objectives are simultaneously achieved. Gray areas in all panels indicate predator extinction. Trophic control parameters set to default values.

### Supplement 5: Effects of trophic forces variation on fishing strategies and objectives conciliation

**Figure S5.**
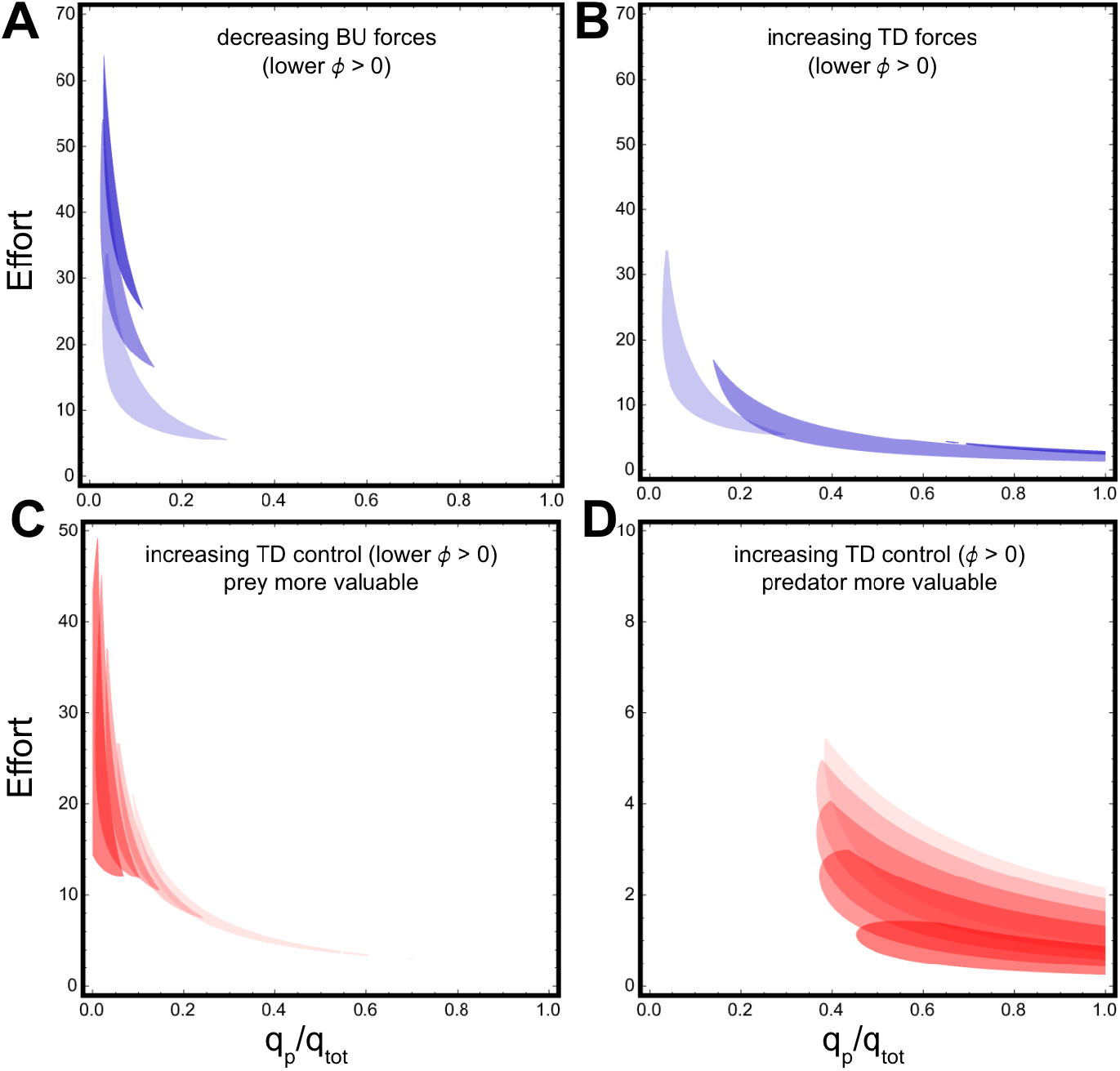
Decreasing bottom-up or increasing top-down forces have contrasting effects on reconciling economic and ecological objectives. The areas displayed correspond to fisheries producing more than 60% of both highest Shannon index value and highest yield. Darker blue shades indicate reduced intraspecific competition among prey (A, weaker bottom-up forces) or increased predation rates by predators (B, stronger top-down forces). A-B: s_pk_ = 6, s_ck_ = 5. C-D: Mean conciliating area for different parameters combinations producing a given *ϕ* value. Lighter red shades correspond to lower *ϕ*, as more top-down controlled systems.

### Supplement 6: Price asymmetries control most profitable fishing strategies and economic-ecological trade-offs (low BU control)

**Figure S6.**
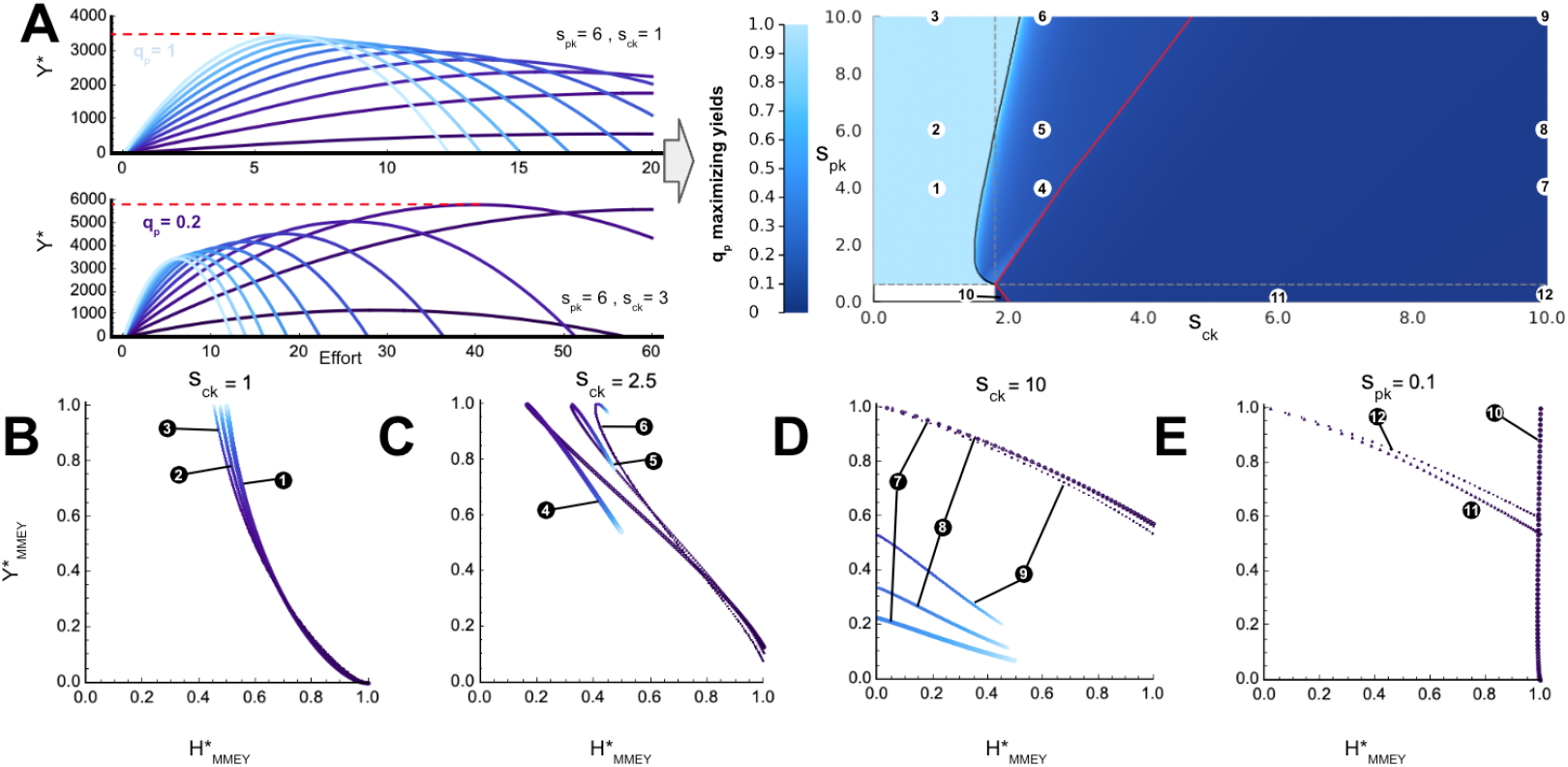
Impact of price structure (*s*_*ck*_, *s*_*pk*_) on strategies maximizing yield (A) and associated trade-offs (B-C-D-E). In contrast to Fig. 3, the trophic control is less BU-oriented (*c*_0_ = 0. 3). For a given price structure, the strategy (*q*_*p*_ /*q*_*tot*_) which maximizes profits among all possible strategies is depicted on panel A (right). Dashed lines: price structures for which monospecific predator (horizontal) or prey (vertical) fishing is unprofitable. Black line: prey yield contribution is significant compared to predator. Red-bordered area: price structures for which maximizing profits leads to predator extinction. B-C-D-E: Impacts of optimizing economic yields on 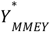 and on the resulting 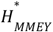 (normalized). Dot colors and numbers indicate the strategy and the associated price structure (A).

### Supplement 7: Price asymmetries control most profitable fishing strategies and economic-ecological trade-offs (TD control)

**Figure S7.**
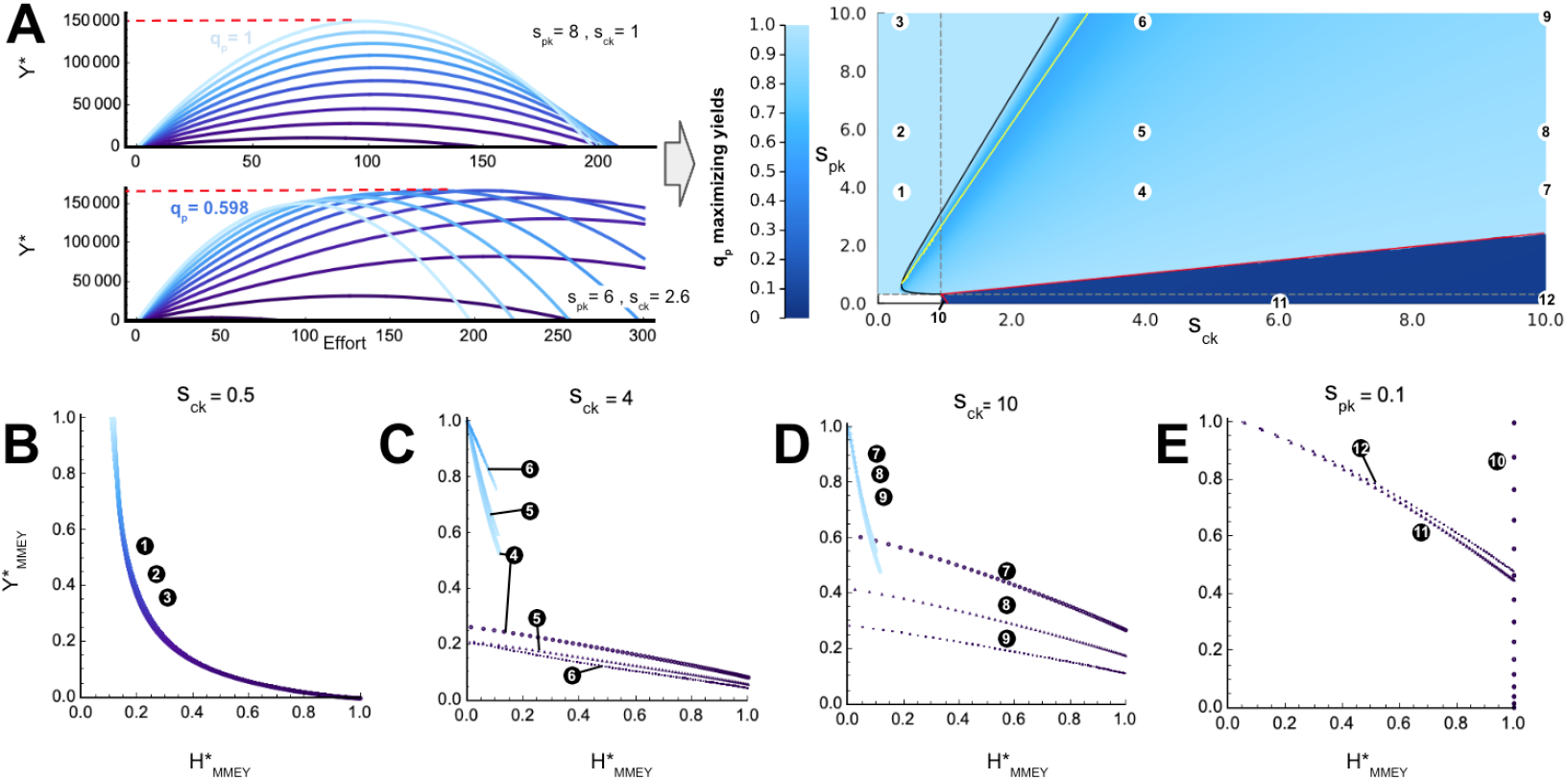
Impact of price structure (*s*_*ck*_, *s*_*pk*_) on strategies maximizing yield (A) and associated trade-offs (B-C-D-E). In contrast to Fig. 3, the trophic control is TD (*ϕ* < 0, *c*_0_ = 0. 02). For a given price structure, the strategy (*q*_*p*_ /*q*_*tot*_) which maximizes profits among all possible strategies is depicted on panel A (right). Dashed lines: price structures for which monospecific predator (horizontal) or prey (vertical) fishing is unprofitable. Black line: prey yield contribution is significant compared to predator. Red-bordered area: price structures for which maximizing profits leads to predator extinction. B-C-D-E: Impacts of optimizing economic yields on 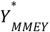 and on the resulting 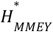 (normalized). Dot colors and numbers indicate the strategy and the associated price structure (A).

### Supplement 8: Maximizing profits makes fisheries economic and ecological status more vulnerable to errors (low BU control)

**Figure S8.**
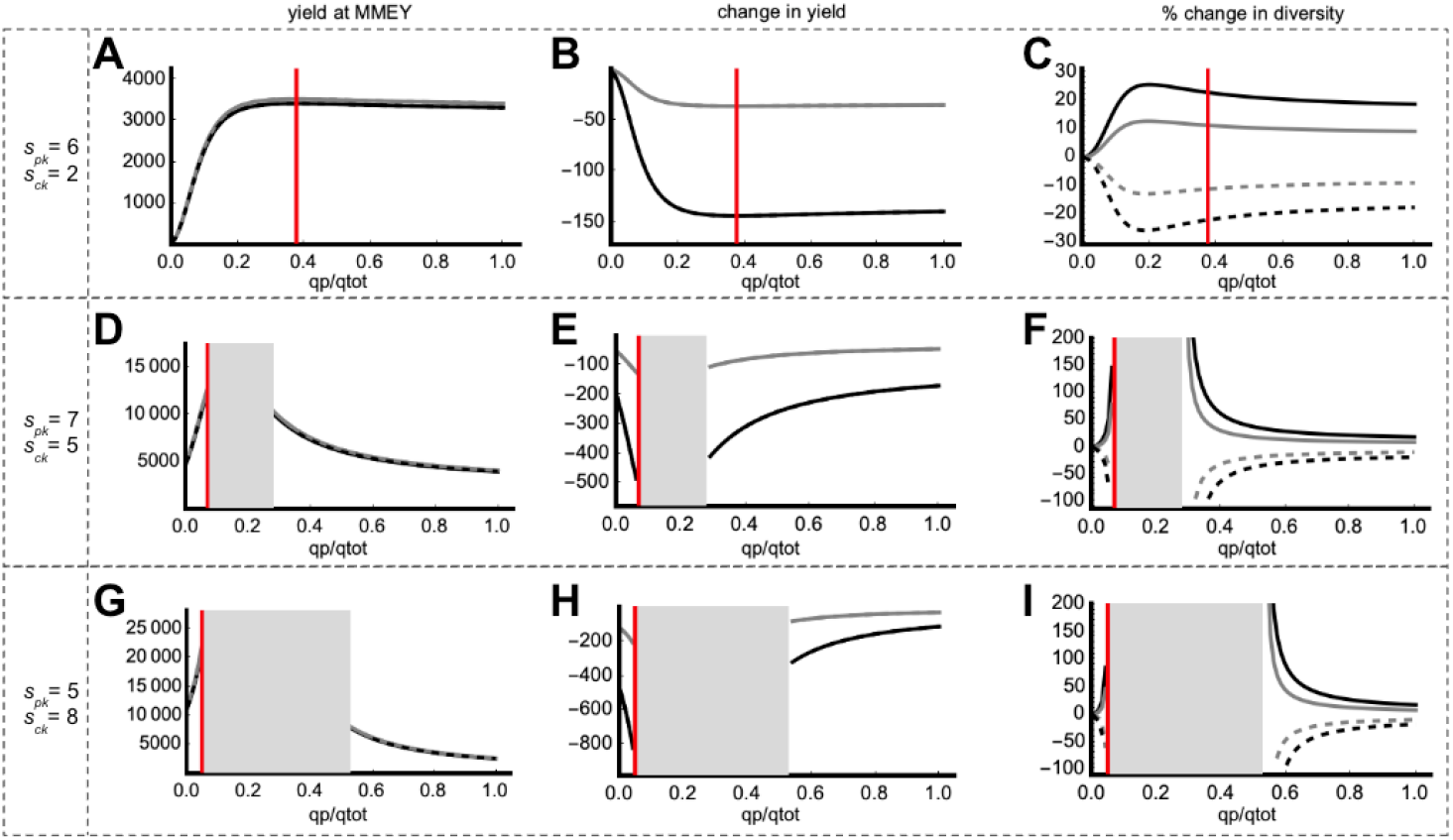
Impact of estimation errors in *E*_*MMEY*_ calibration on economic yield and diversity. Black (gray) lines: error of 20% (10%) in *E*_*MMEY*_ calibration. Solid (dashed) lines: underestimation (overestimation). The red vertical line depicts the fishing strategy producing the best economic yield. A-D-G: economic yields with calibration errors. B-E-H: effects of calibration errors on yields. C-F-I: effects of calibration errors on diversity. Price structure values are indicated on the left side of each line. Black areas: unprofitable fishing strategies. Gray areas: fishing strategies leading to predator extinction when maximizing yields with MMEY metrics (calculated without calibration errors). The system is less bottom-up (more top-down) controlled compared to Fig. 4 (*ϕ* > 0, *c*_0_ = 0. 3).

### Supplement 9: Maximizing profits makes fisheries economic and ecological status more vulnerable to errors (TD control)

**Figure S9.**
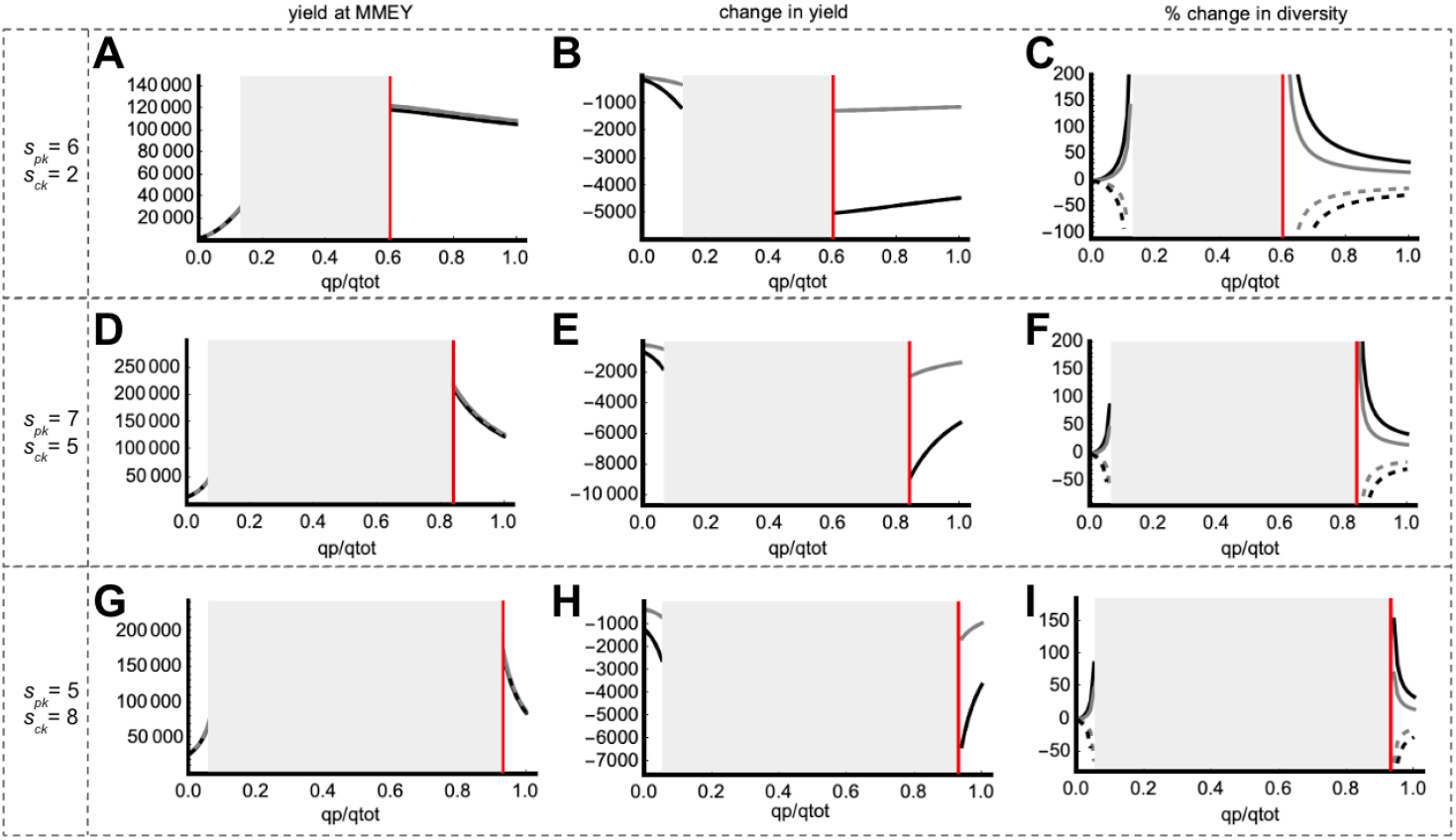
Impact of estimation errors in *E*_*MMEY*_ calibration on economic yield and diversity. Black (gray) lines: error of 20% (10%) in *E*_*MMEY*_ calibration. Solid (dashed) lines: underestimation (overestimation). The red vertical line depicts the fishing strategy producing the best economic yield. A-D-G: economic yields with calibration errors. B-E-H: effects of calibration errors on yields. C-F-I: effects of calibration errors on diversity. Price structure values are indicated on the left side of each line. Black areas: unprofitable fishing strategies. Gray areas: fishing strategies leading to predator extinction when maximizing yields with MMEY metrics (calculated without calibration errors). The system is top-down controlled (*ϕ* < 0, *c*_0_ = 0. 02).

### Supplement 10: Mathematical developments using MMEY metrics

In an exploited predator-prey system, we assume that fishers:

- face greater constraints in modifying the technology outfitting their vessels than in adjusting the number of days at sea (effort is more readily variable than technology, and thus fishing strategy),
- they adjust the effort to maximize their incomes (yield, see eq. 9) given the technology level *q*_*p*_ they have at their disposal and the prevailing price structure for fish sales (*s*_*pk*_, *s*_*ck*_).

We aim to understand how price conditions influence optimal fishing from a yield perspective. Prey (*C*) and predators (*P*) interact according to:

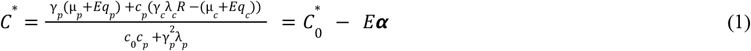

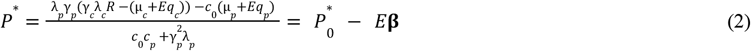

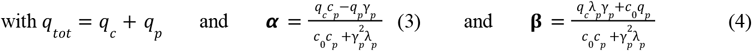

*s*_*p*_ and *s*_*c*_ represent the price of one unit of predator and one unit of prey, respectively. Assuming a linear cost ε associated with effort, the profitability of a fishery is assessed by its economic yield at effort *E*, given by:

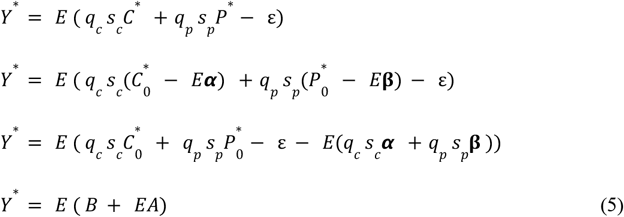

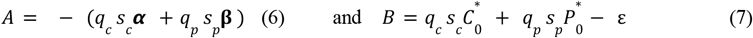

We observe that for *Y*\* to be positive over 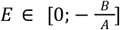,it is necessary that *A* < 0 (a constraint on the parabolic shape) and thus *B* > 0 (since *E* is defined as positive). Fishers maximize economic returns when fishing at *E*_*MMEY*_ :

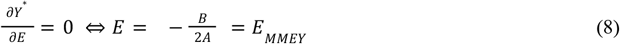

Fishers then achieve 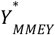 revenues:

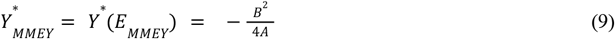

Note that we always consider a coexistence scenario where prey and predators persist together:

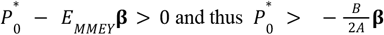

Understanding how price structure affects the most profitable fishing strategy requires examining the sign of 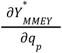.Indeed, if

- 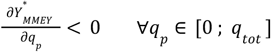 then the most profitable allocation of effort is the one focused solely on targeting preys.
- 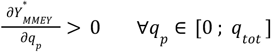 then the most profitable allocation of effort is the one focused solely on targeting predators.
- 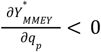 then 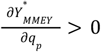 with *q*_*p*_ ∈ [0 ; *q*_*tot*_] then the most profitable allocation of effort is a mixed distribution, where preys and predators are both targeted.

It turns out that the maximum economic yield varies with the fishing strategy according to:

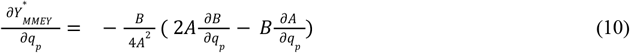

Thus, to determine the sign of 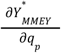,we will look for:

1. the major fishing regimes as a function of the price structure
2. the most profitable fishing strategy: theoretical scenarios
3. demonstrate why:
  a. using an MMEY metric, the most profitable fisheries are those targeting only predators when predators are the most valuable
  b. market conditions shift the situation from one focused on predators to a situation where the most profitable fishing strategy is a mixed approach
  c. the relative impacts of errors in assessment or implementation of economic yields are not dependent from fishing strategy

#### 1) Major fishing regimes as a function of the price structure

From eq. 7 and eq. 5 it is observed that: 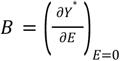

B represents the slope of the yield when evaluated at the origin. B > 0 means that, given the available technologies (strategy *q*_*p*_), a situation with fishing is more advantageous than a situation without fishing (it generates yield, even for an effort that approaches 0). Conversely, B < 0 means that, with the technologies available to the fishers, going to sea is always disadvantageous compared to a situation where they do not fish (fishing is always unprofitable, regardless of the effort). From the perspective of using MMEY as a management tool, B > 0 means that *E*_*MMEY*_ > 0, and thus the maximum yield is positive and achieved at *E*_*MMEY*_ . When B < 0, *E*_*MMEY*_ < 0, and thus the best yield is zero. Therefore, for fishing to be financially viable and for MMEY to be used in management, B must be positive.

It can be noted that *B* can be rewritten in the form:

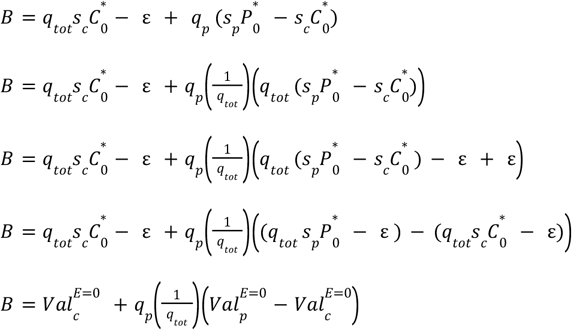

With 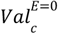 and 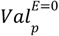,the valuation of the prey stock and the predator stock without fishing, respectively.

Two cases can be observed:

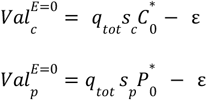

- if 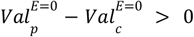 the valuation of predator stocks is higher than that of prey stocks. In this case, fishing is possible as long as the technologies allocate a certain portion of the total effort towards predators:

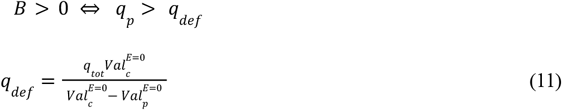

With *q*_*def*_ the threshold strategy for which *B* = 0. This catch of predators can indeed be zero (*q*_*def*_ < 0 when 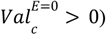, which means that all strategies generate yield (*q*_*p*_ ≥ 0 > *q*_*def*_) as long as the fishers are fishing.

However, since *q*_*p*_ > *q*_*def*_ to allow fishing, *q*_*def*_ must be lower than *q*_*tot*_ . If *q*_*def*_ > *q*_*tot*_, it means that all the strategies available to the fishers always produce negative yields. Therefore, for *q*_*def*_ < *q*_*tot*_, it must be the case that:

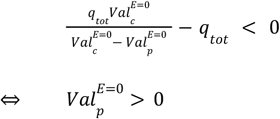

In other words, for there to be at least one effort distribution that generates yield, it is necessary that the predator stock has a positive valuation. This situation is therefore summarized under the term “*fishing allowed by predator valuation*”.

- if 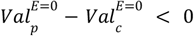 when the valuation of predator stocks is lower than that of prey stocks, fishing is possible as long as the technologies allocate a certain portion of the total effort towards the prey:

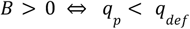

This catch can indeed be zero (if *q*_*def*_ > *q*_*tot*_), which means that all strategies generate yield (*q*_*p*_ ≤ *q*_*tot*_ < *q*_*def*_) as long as the fishers are fishing.

However, since *q*_*p*_ < *q*_*def*_ in order to fish, *q*_*def*_ must be positive. If *q*_*def*_ < 0, it means that all the strategies available to the fishers always produce negative yields. Therefore, for *q*_*def*_ > 0, it must be the case that:

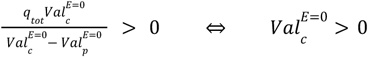

In other words, for there to be at least one distribution of effort that generates a positive yield, it is necessary that the prey stock has a positive valuation. We summarize this situation under the term “*fishing authorized by prey valuation*”.

Thus, there are two main fishing regimes: a “*fishing authorized by predator valuation*” regime, when the valuation of predator stocks is higher than that of prey stocks, and a “*fishing authorized by prey valuation*” regime in the opposite situation.

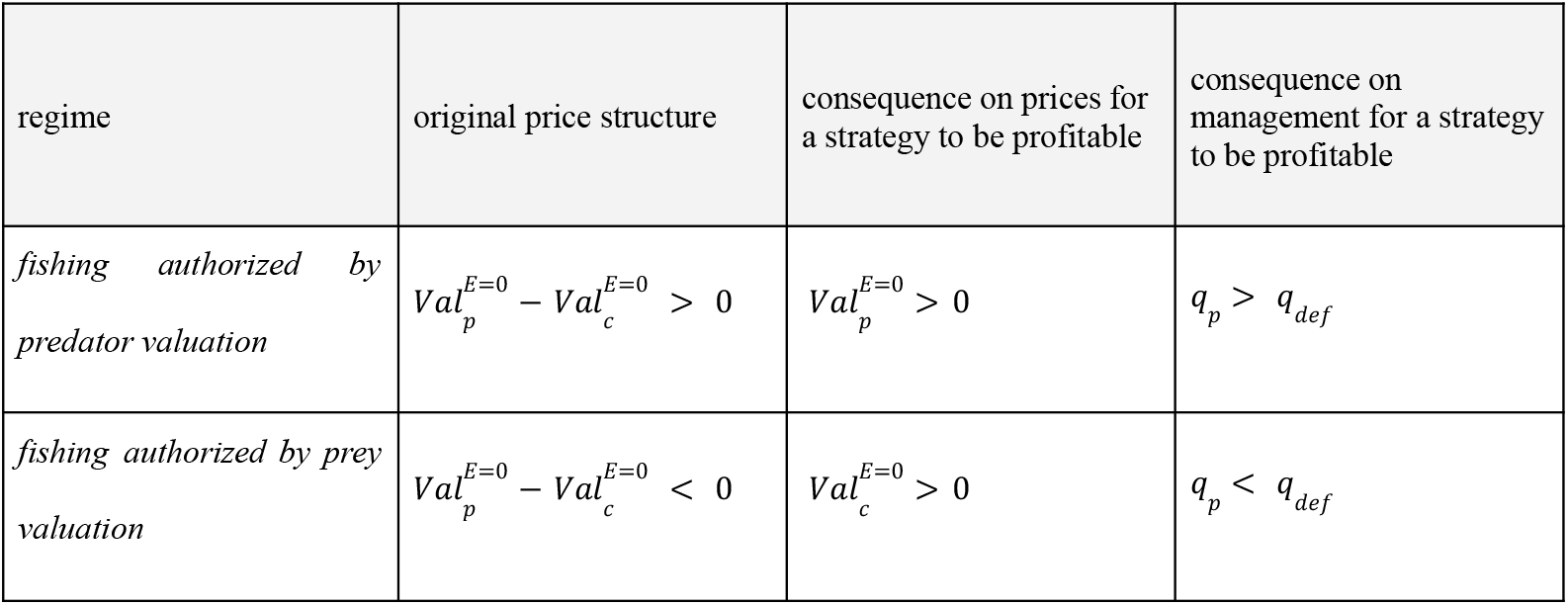

Outside of these conditions, it is materially impossible to fish.

#### 2) Most profitable fishing strategy: theoretical scenarios

From eq. 10, we can find the sign of 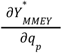 based on that of 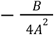 and 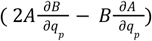.Recall that fishing is only feasible if B > 0 and A < 0 .

<u>Sign of</u> 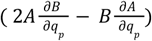:

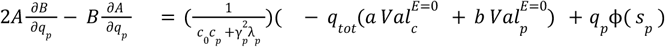

with

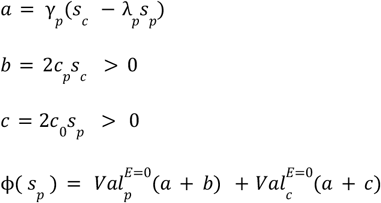

thus 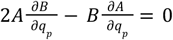 when *q*_*p*_ = *q*_*def*:_

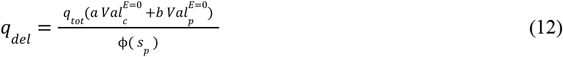

To find the sign of 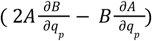,we need to understand its monotonicity.

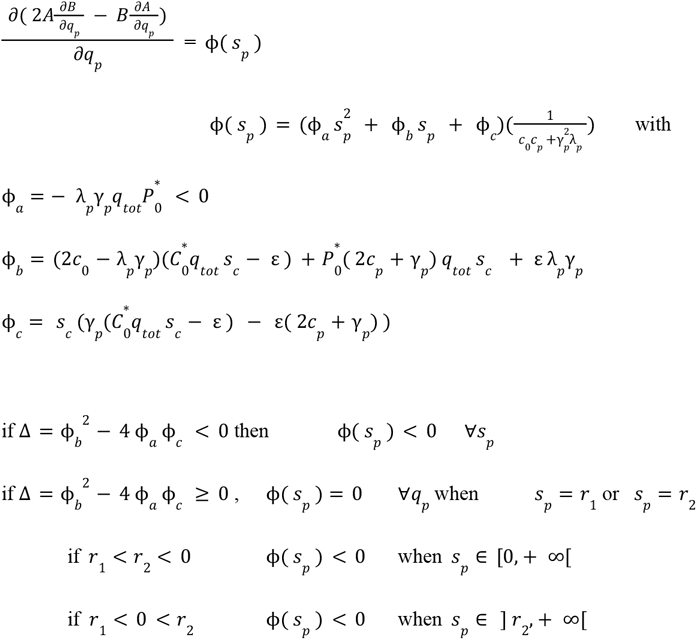

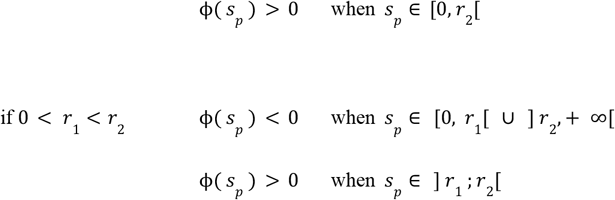

We will subsequently assume the strict monotonicity of *r*_1_ and *r*_2_ over intervals where they are continuous.

In the case of the regime of “*fishing authorized by predator valuation*”, we obtain the following theoretical cases:

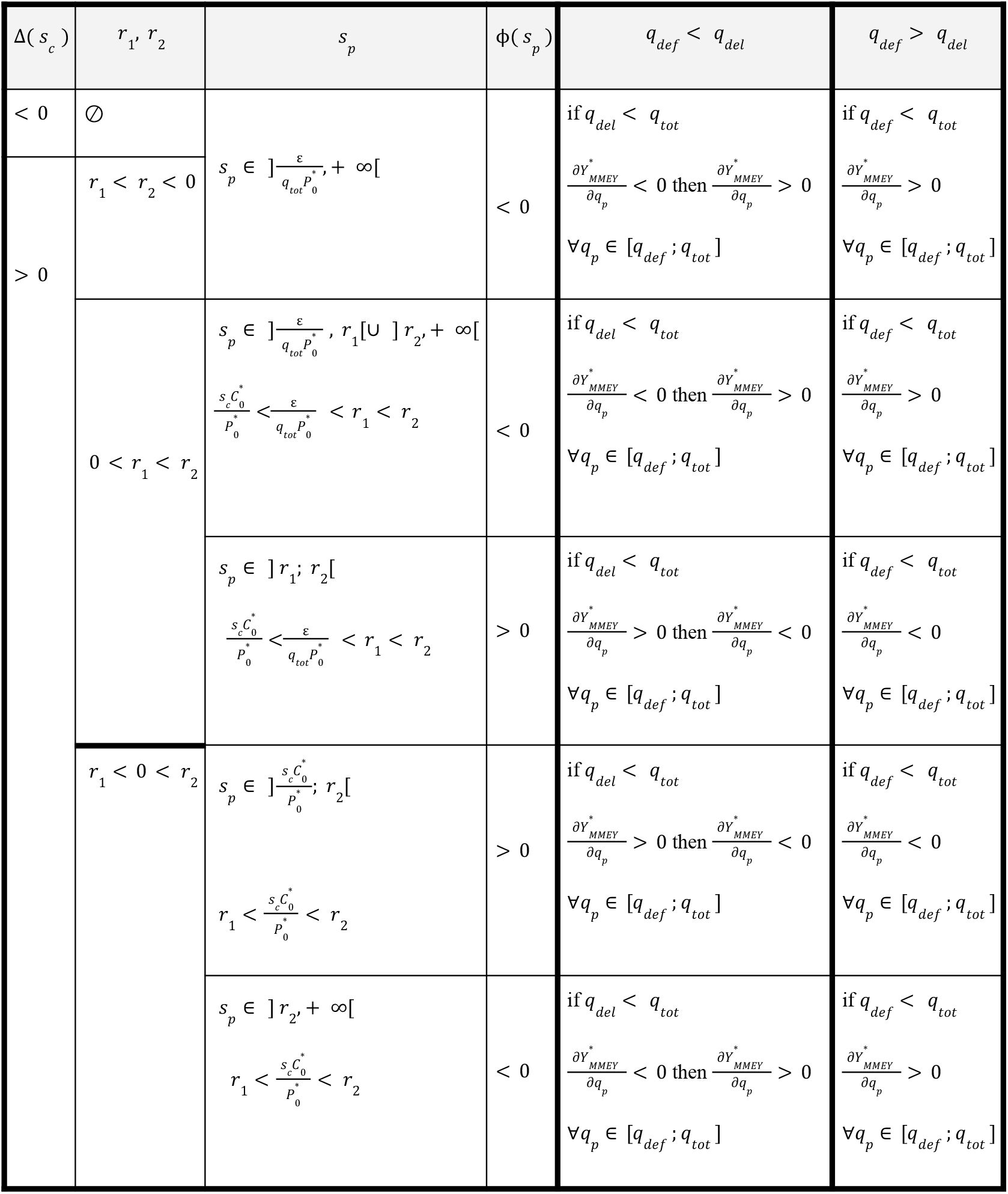

#### 3) a) Using an MMEY metric, the most profitable fisheries are those targeting only predators when predators are the most valuable

We consider the case where Δ(*s*_*c*_) > 0 and *r*_1_ < *r*_2_< 0. First, we examine the conditions on *s*_*c*_ needed to satisfy these inequalities.

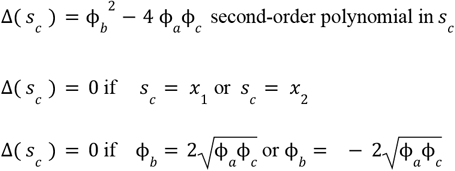

We then seek the sign of ϕ_*b*_ (*x*_1_):

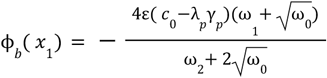

with

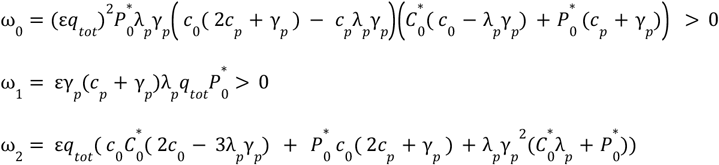

Thus, when *c*_0_ − *λ*_*p*_ *γ*_*p*_ > 0, ϕ_*b*_ (*x*_1_) can be positive only when

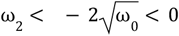

but when *c*_0_ = 0 (most extreme case),

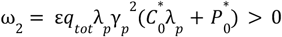

Thus

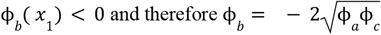

and thus 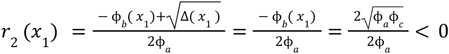 because ϕ_*a*_ < 0 ∀ *s*_*c*_

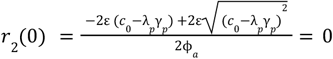

In this way, for ∀*s*_*c*_ ∈ [0 ; *x* _1_], *r*_1_ < *r*_1_ < 0

Similarly, when

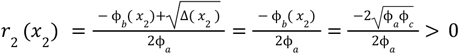

and

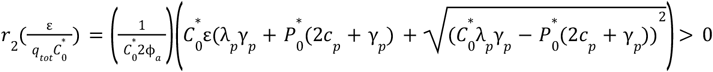

Thus 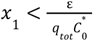 (and therefore 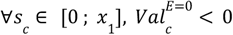).

Then, 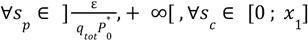

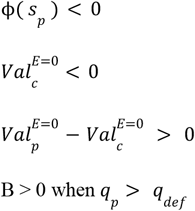

And because of eq. 10

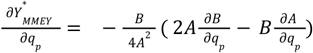

We seek to determine whether *q*_*del*_ > *q*_*del*_ or *q*_*del*_ > *q*_*del*_ in order to understand how 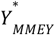 varies when moving the *q*_*p*_ > *q*_*del*_ . Specifically, in the first case, when ∀*q*_*p*_ ∈ [*q*_*del*_, *q*_*tot*_], the yield at MMEY would be increasing and then decreasing with the fishing strategy, whereas in the second case, ∀_*p*_ ∈ [*q*_*del*_, *q*_*tot*_], the yield at MMEY would be increasing with the fishing strategy.

Thus,

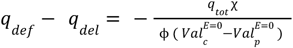

With

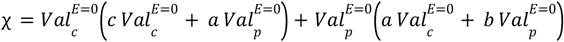

and because

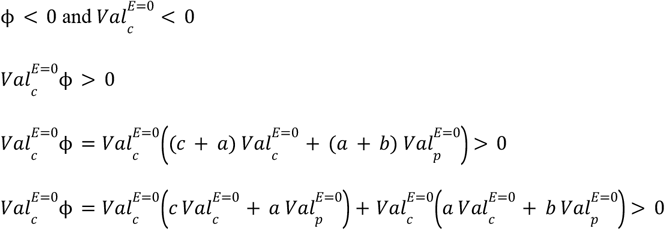

and

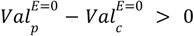

thus

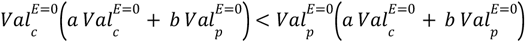

and therefore

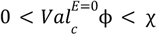

We conclude that under these conditions, χ > 0 and thus *q*_*def*_ − *q*_*del*_ < 0

Hence, 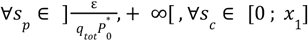

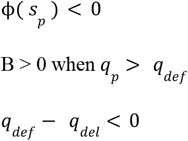

Therefore, fishing is only feasible when *q*_*p*_ ∈ [*q*_*def*_ ; *q*_*tot*_] and within this range 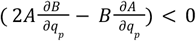.

Consequently, 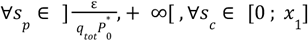

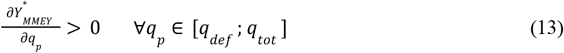

In these conditions, the most profitable fishing strategy using MMEY metrics is one where all the effort is allocated to the capture of predators.

#### 3) b) Market conditions that shift from pure predator fishing to mixed fishing

In the case where *s*_*p*_ ∈] *r*_1_ ; *r*_2_ [and 0 < *r*_1_ < *r*_2_, the most profitable allocation of effort over the interval [*Max*(*q*_*def*_, 0) ; *q*_*tot*_] is a mixed allocation if 0 < *q*_*del*_ < *q*_*tot*_ . Indeed, when *q*_*del*_ ≥ *q*_*tot*_, 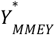 remains increasing over the interval [*Max*(*q*_*def*_, 0) ; *q*_*tot*_], and thus, the most profitable allocation is an allocation where *q*_*p*_ = *q*_*tot*_, meaning an allocation where only predators are targeted for fishing. Conversely, when 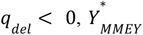 is decreasing over the interval [*Max*(*q*_*def*_, 0) ; *q*_*tot*_], and thus the most profitable allocation is an allocation where *q*_*p*_ = 0, meaning an allocation where only prey are targeted for fishing.

Thus, ∀*s*_*p*_ ∈] *r*_1_ ; *r*_2_ [and ∀*s*_*c*_ such that 0 < *r*_1_ < *r*_2_,

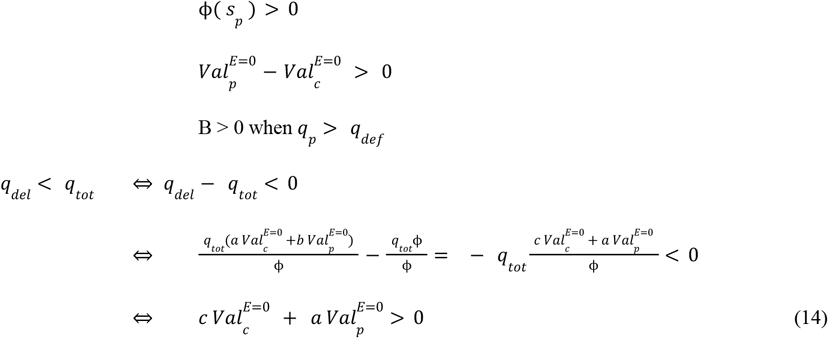

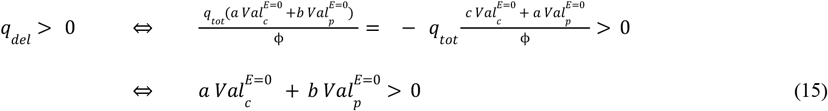

Thus, in the space *s*_*p*_ ∈] *r*_1_ ; *r*_2_ [and *s*_*c*_ such that 0 < *r*_1_ < *r*_2_, there exists a subspace (*s*_*c*_, *s*_*p*_) in which equations 14 and 15 are satisfied.

Recall that:

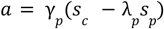

When fishers target only predators, this indirectly affects the prey density positively according to 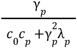 (eq. 1), and thus affects the valuation of prey as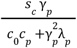.

When fishers target only prey, this indirectly affects the predator density negatively according to 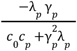 (eq. 2), and thus affects the valuation of predators as 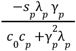.

Thus, targeting only predators means missing out on a valuation gain from the prey 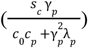,but not being penalized by the lower productivity of the predators 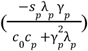.

Thus, *a* represents the foregone gain from fishing only predators. It can be positive (real foregone gain from targeting only predators) or negative (gain realized).

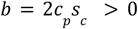

When fishers target only prey, this results in a direct decrease in their density according to 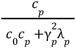 (eq. 1), and consequently affects their valuation according to 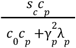.

Thus, *b* represents the direct foregone gain in the valuation of prey due to targeting prey.

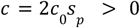

When fishers target only predators, this results in a direct decrease in their density due to 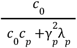. (eq. 2), and consequently, in the valuation of predators due to 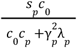.

Thus, *c* represents the direct foregone gain in the valuation of predators due to targeting predators.

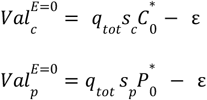

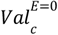 and 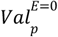 represent the valuation of the prey and predator stocks, respectively, in the absence of fishing. These values reflect the marginal gain from fishing for prey or predators.

We seek to understand what represents (eq. 14): 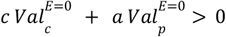

Thus, targeting only predators results in an intrinsic foregone gain (*c*) on the valuation of their stock and missing the exploitation of the prey stock 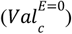.

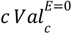 then represents the direct foregone gain of not targeting prey.

Targeting only predators also implies an indirect foregone gain of not exploiting a more highly valued prey stock 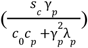,but without suffering from lower productivity 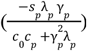.

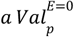 represents the indirect opportunity cost (due to the feedbacks between prey and predators) of not fishing for prey.

Thus, when 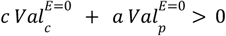,this means that the total foregone gain (direct + indirect) is positive, in other words, the most profitable strategy is not to focus solely on fishing predators.

We seek now to understand what represents (eq. 15): 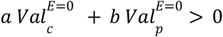

Fishing exclusively for prey means having a direct intrinsic foregone gain (*b*) on the valuation of their stock and missing out on exploiting the predator stock 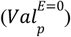

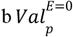 then represents the direct foregone gain of not fishing predators.

Fishing exclusively for prey also means having an indirect foregone gain by not exploiting a more highly valued predator stock 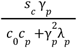, but without suffering from the reduced productivity 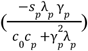 of the predators, as they are not being fished.

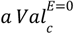 then represents the indirect opportunity cost (through feedbacks between prey and predators) of not fishing predators.

Thus, when 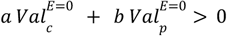,this means that the total foregone gain (direct + indirect) is positive, in other words, the most profitable strategy is not to fish exclusively for prey

Thus, it is clear that when both conditions are met, there is an foregone gain to fishing only for prey 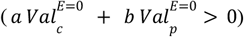 but also an foregone gain to fishing only for predators 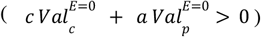. The most profitable strategy is then necessarily a mixed strategy 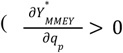 followed by 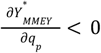 when *q*_*p*_ ∈ [*Max*(*q*_*def*_, 0) ; *q*_*tot*_]).

#### 3) c) The relative impacts of errors in assessment or implementation on economic yields are not dependent from fishing strategy

When fishermen use MMEY metrics to maximize their economic yield, they have to fish with an effort of 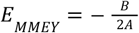 (eq. 8). Let’s assume they’re wrong by as much as ***θ*** (positive or negative) in *E*_*MMEY*_ assessment. It induces:

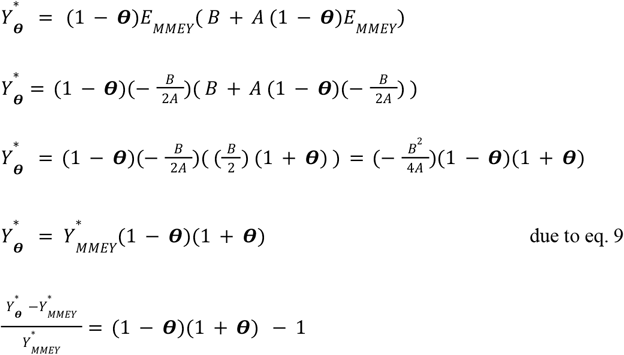

Therefore, the relative change in economic yields does not depend on the fishing strategy (*q*_*p*_).

